# Global metagenomics reveals plastid diversity and unexplored algal lineages

**DOI:** 10.1101/2025.03.28.644651

**Authors:** Bikash Shrestha, Miguel F. Romero, Juan C. Villada, Plastid MAG Consortium, Blaby-Haas Crysten, Frederik Schulz

## Abstract

Photosynthetic organelles in eukaryotes originated through primary endosymbiosis with a cyanobacterium, an event that profoundly shaped the evolutionary landscape of the eukaryotic tree of life. Primary plastids in Archaeplastida, especially in cultivable plants and algae, contribute most to known plastid diversity. Secondary and higher-order endosymbiosis, involving eukaryotic hosts and algal endosymbionts, further spread photosynthesis among protists within the CASH lineages (Cryptophyta, Alveolata, Stramenopila, and Haptophyta). Despite various hypotheses explaining secondary plastid evolution and distribution, empirical support remains limited. Here, we employ cultivation-independent global metagenomics to expand plastid diversity and investigate plastid origins. We captured 1,027 plastid sequences, including 300 novel sequences belonging to previously unsequenced plastids and representing yet-to-be described microeukaryotes. This includes a new lineage that offers insights into plastid evolution in haptophytes and cryptophytes. Our results confirm that Archaeplastida plastids originated from an early-branching cyanobacterial lineage closely related to Gloeomargaritales and identify the closest extant relative of *Paulinella* plastids. Additionally, our findings suggest two independent origins of secondary red algal plastids, contributing to plastid diversity in CASH lineages and challenging the prevailing model of single secondary plastid origin. Our study highlights the importance of metagenomic data in uncovering biological diversity and advancing understanding of plastid relationships across photosynthetic eukaryotes.

## INTRODUCTION

Oxygenic photosynthesis is a crucial metabolic process that has profoundly shaped life on earth. While oxygenic photosynthesis (referred herein as photosynthesis) first evolved in cyanobacteria, autotrophy has spread between kingdoms, from bacteria to eukaryotes, and across Eukarya through endosymbiosis, leading to the emergence of taxonomically distinct photosynthetic eukaryotes. The resulting plants and algae play essential roles in terrestrial and aquatic ecosystems, with plastids serving as their primary organelles responsible for generating molecular oxygen and converting CO_2_ to sugars.

Plastids originated through primary endosymbiosis, where a heterotrophic eukaryote engulfed a now extinct cyanobacterium that eventually evolved into a photosynthetic organelle. To date, only two confirmed instances of primary endosymbiosis exist. The first event, occurring approximately 1.5 billion years ago, involved a eukaryote and β-cyanobacterium endosymbiont, leading to the formation of plastids in Archaeplastida, a group that includes green algae and land plants (Viridiplantae), red algae (Rhodophyta), and glaucophytes (Glaucophyta)^1–5^. Despite some uncertainty about the cyanobacterial ancestor of these plastids, recent research indicates that the ancestor is related to early-diverging lineages, such as Gloeomargaritales^6,7^. The monophyly of Archaeplastida plastids is generally well-supported by conserved gene content, synteny, presence of inverted repeats, and high sequence similarity of 16S rRNA genes^8,9^. However, studies challenging this monophyly are not uncommon and phylogenetic relationships among the Archaeplastida group remain debated^8–10^. The second primary endosymbiosis took place much more recently, between 90-140 million years ago involving *Paulinella* (Rhizaria), an amoeba that acquired an α-cyanobacterium related to the *Synechococcus*/*Prochlorococcus* clade^11–13^. Unlike Archaeplastida plastids, the *Paulinella* plastid retains more of its cyanobacterial genome (∼1 Mb vs. <200 Kb) and has distinct membrane structure^14^.

Beyond primary endosymbiosis, the genesis of new algal lineages has occurred through secondary or serial endosymbiosis, where non-photosynthetic eukaryotes have engulfed an alga forming secondary/complex plastids^9,10^. Algae such as Chlorarachniophytes, Euglenophyceae, and dinoflagellates acquired secondary plastids through the endosymbiosis of distinct green algae, and the evolutionary origins of these independent events are relatively well understood^15^. However, the evolutionary origins and spread of red alga-derived plastids in algal groups, including Cryptophyta, Haptophyta, Ochrophyta (photosynthetic Stramenopila), and Alveolates (comprising Colpodellida, Apicomplexa, and dinoflagellates), collectively known as CASH lineages, remain contentious. Several hypotheses have been proposed to account for the presence of red algal-derived plastids in distinct lineages of protists. The Chromalveolate hypothesis, first proposed by Cavalier-Smith^16^, suggests that the CASH lineages share a common algal ancestor, with multiple plastid losses to explain non-photosynthetic protist lineages. However, phylogenomic studies imply that multiple plastid losses would be highly improbable^17,18^. An alternative model, the Rhodoplex hypothesis, proposes that secondary/complex red algal plastids originated within one CASH lineage (likely Cryptophyta) and subsequently spread via tertiary or higher order endosymbiosis (serial endosymbiosis^18–21^). Despite these competing hypotheses, empirical evidence supporting them remains limited^22^.

Our understanding of plastid evolution has primarily relied on complete plastid genomes (plastomes) obtained from isolates or culture collections. As a result, currently available plastid sequences likely represent only a fraction of existing diversity, as many environments remain underexplored. The use of metagenome-assembled genomes (MAGs) has significantly expanded our understanding of microbial diversity, since sampled organisms do not need to be isolated or cultured. Research leveraging MAGs has been instrumental in discovering new lineages across the major domains of life and elucidating their evolutionary history and relationships^23–25^. The analysis of plastid MAGs (ptMAGs) has the potential to uncover previously unidentified plastid lineages, suggest the existence of new algal lineages, and offer deeper insight in our current understanding of plastid evolution.

Here, we aimed to investigate plastid diversity and evolution using metagenomic approaches. We generated a dataset of ptMAGs and expanded the Cyanobacteriota MAGs dataset using publicly available metagenomic data in the Integrated Microbial Genomes and Microbiomes (IMG/M) database^26,27^. By integrating these MAGs with reference plastomes and Cyanobacteriota genomes, we reconstructed plastid evolutionary relationships. Our analyses reveal the closest cyanobacterial relatives of *Paulinella* plastids, confirm Archaeplastida plastids originated from an early-branching cyanobacterial lineage, and provide evidence for two independent events involving engulfment of a red alga. Additionally, we report multiple plastid lineages that likely represent previously undiscovered algal lineages and provide complete or nearly complete genomic sequences for plastids previously identified solely through rRNA sampling, including a novel plastid lineage that offers new insights into plastid evolution within haptophytes and cryptophytes.

## MATERIALS AND METHODS

### Phylogenetic markers

We utilized two sets of phylogenetic markers, referred to as UNI56 and PLASTID54, for all our phylogenetic analyses (Supplementary Table S1). The UNI56 markers included 56 universal single-copy marker proteins, comprising 30 large and small ribosomal subunits, three DNA-directed RNA polymerase subunits,10 tRNA synthetase, and other functional proteins^28,29^. The PLASTID54 consisted of 34 Hidden Markov Model (HMM) profiles derived from the UNI56 markers, including large and small ribosomal subunits, DNA-directed RNA polymerase subunits, and the preprotein translocase SecY. Additionally, they included 20 HMM profiles for plastid photosynthetic proteins associated with photosystems, ATP synthase, and the cytochrome complex as available in Pfam-A HMMs (last accessed August 2024) and as detailed in Supplementary Table S1. The phylogenomic relationships among Cyanobacteriota species were inferred using the UNI56 markers, while the plastid phylogeny among photosynthetic eukaryotes was estimated using the PLASTID54 markers. The initial placement of plastids within the cyanobacterial lineage was examined using the UNI56 markers and then re-evaluated with the PLASTID54 markers.

### Cyanobacteriota genomes and MAGs

We performed metagenomic binning on 31,152 metagenomes from the IMG/M^26,27^, focusing on non-redundant samples publicly available up to April 2023^30–38^. Only contigs with lengths of ≥ 5 Kb were processed. The binning was conducted on each sample using MetaBAT2 with parameters: --minContig 5000, --minClsSize 20000, and –cvExt, and the gene calling on each bin was performed using Prodigal (v2.6.3)^39^.

Reference genomes and bins from publicly accessible repositories were downloaded from the IMG isolates repository (n = 115,969) and the bacterial and archaeal representative datasets of GTDB (n = 85,205)^40^. All genomes and bins underwent CheckM v1.1.3^41^ analysis using UNI56 markers for universal marker detection required for phylogenomic reconstruction, CheckM2 for quality assessment, and seqkit v2.5.1^42^ for assembly characterization. Genomes and bins with less than 50% UNI56 markers, CheckM2 completeness below 50%, or CheckM2 contamination above 5% were excluded from the genome catalog.

Species-level clusters were inferred with skani (v0.1.4^43^, parameters: ANI ≥ 95%, align_fraction_ref ≥ 50%, and align_fraction_query ≥ 50%) and cluster representatives were selected based on the highest quality metrics (completeness and contamination), with ties resolved by selecting assemblies with the highest number of predicted genes and the highest N50 value. Finally, genomes classified as Cyanobacteriota according to the GTDB taxonomic classification were selected for the downstream analyses.

### Cyanobacteriota phylogeny and diversity

The phylogenetic relationship of Cyanobacteriota was inferred with an estimated species tree using the nsgtree pipeline (https://github.com/NeLLi-team/nsgtree) and IQ-TREE v.2.0^44^. In brief, the nsgtree pipeline identified orthologs associated with UNI56 markers using hmmsearch v. 3.1/b2 (www.hmmer.org). For each genome and marker, the ortholog with the highest bitscore was extracted. Extracted orthologs were then aligned using MAFFT v.7^45^ and trimmed using trimAl v.1.4^46^ with a gap threshold of 10% (−gt 0.1). A supermatrix alignment was generated by concatenating 56 individual ortholog alignments. Only genomes retaining at least 50% of the UNI56 markers were included in the final supermatrix alignment. A maximum-likelihood (ML) species tree was estimated with the supermatrix alignment using IQ-TREE with the LG+F+I+G4 substitution model, and branch support was evaluated with 1000 ultrafast bootstrap pseudoreplications^47^. The percentage increase in phylogenetic diversity was calculated by comparing the sum of branch lengths of the Cyanobacteriota species trees with and without new bacterial MAGs generated in our study (labeled as NeLLi2023 in Supplemental Table S2).

To generate representative Cyanobacteriota genomes for examining the origin of plastids, we further dereplicated the Cyanobacteriota dataset. Pairwise evolutionary distances (branch lengths) were inferred between all species-level Cyanobacteriota culture representatives utilizing the ML species tree generated with IQ-TREE using PhyloDM^48^ and then clustered with MCL at different distances cutoff values, ranging from 0.2 to 1.0 and an inflation value at 1.5^49^. For each cutoff value, we calculated basic cluster statistics, such as, the number of clusters, number of singletons, average cluster size, and the number of genomes in the largest cluster, and sorted the cluster based on the UNI56 marker count for the genomes/MAGs generated by the nsgtree pipeline. We inspected the clusters at different cutoffs to ensure the genomes belonging to different GTDB taxonomic orders were not collapsed or mixed within clusters, allowing us to select an appropriate cutoff value.

### Generation of plastid metagenomes

We analyzed 26,257 public metagenomes from the IMG/M database to identify contigs containing plastid 16S rRNA gene sequences using cmsearch (Infernal v1.1)^50^ with the bacterial SSU rRNA covariance model RF00177 from the Rfam database. To accommodate longer sequences, we utilized the --anytrunc option in cmsearch and parsed the output to identify contigs with sequential and multiple non-overlapping alignments to the covariance model, which were then assembled into single sequences. The recovered SSU rRNA sequences were annotated using the SILVA^51^ and PR2^52^ databases via blastn v.2.13.0^53^, retaining those with a plastid sequence as the best hit and an alignment length of at least 500 bp, resulting in 11,298 candidate plastid SSU rRNA genes. For further analyses, we selected 4,497 contigs with lengths of at least 5 Kb.

### Filtering plastid metagenomes

To identify high-quality ptMAGs for the study, we performed structural gene annotation using Prodigal v2.6.3 with the “-p meta” option, which employs precalculated training files, and keeping other parameters at default settings. We selected ptMAGs that contained at least 10% of UNI56 markers, in accordance with the nsgtree pipeline. To remove non-plastid sequences likely resulting from mis-assembly and mis-binning, we performed a DIAMOND^54^ BLASTp for all predicted proteins in the ptMAGs against the NCBI non-redundant (nr) database (last accessed August 2024), using an e-value cut-off of 1 × 10^−6^ and minimum of 30% of subject and query coverage. We then assigned taxonomic affiliations for the best hits and categorized the BLAST hits based on amino acid percent identity into three groups: low (<70%), medium (≥70% and <90%), and high (≥90%). To streamline the taxonomic affiliation of the proteins, we retained taxonomic ranks at the Domain and Phylum levels, along with the assigned percent identity category. We then assigned taxonomic affiliations to the contigs based on the simplified taxonomic ranks for the majority of the proteins in each contig. Contigs assigned to bacteria with the medium and high percent identities that were not associated with the Cyanobacteriota were discarded. Only contigs assigned to Eukaryota, Cyanobacteriota, and Bacteria from any other phylum but with low percent identities were selected for the further analysis.

Similarly, to remove potentially mis-identified mitochondrial metagenomic contigs, we compared the ptMAGs against 18,222 complete RefSeq mitochondrial genomes available in the NCBI (https://ftp.ncbi.nlm.nih.gov/refseq/release/mitochondrion/; last access October, 2024). For each plastid contig, we generated BLAST hits (blastn, e-value of 1 × 10^−6^) and estimated pairwise identity using skani^34^ with the “skani dist --slow” option, against the mitochondrial RefSeq database. Based on the percent identity and alignment length (95% nucleotide identity and >95% alignment length), we identified and filtered out any contigs that are likely mitochondrial in origin.

### Estimating plastid phylogeny

We inferred plastid phylogeny by calculating a species tree using the nsgtree pipeline (https://github.com/NeLLi-team/nsgtree) and IQ-TREE v.2.0, with adjustments to the dataset and markers. To determine the origin of plastids, we combined a dereplicated Cyanobacteriota dataset with ptMAGs and RefSeq plastomes and inferred phylogenetic relationships using PLASTID54. Our initial ML tree estimation included a dataset containing 259 dereplicated Cyanobacteriota taxa, 236 RefSeq NCBI plastomes, and 647 metagenomic contigs containing plastid 16S rRNA genes. Although we initially had 1,027 MAGs containing 16S rRNA genes, applying the criterion of at least 10% of PLASTID54 markers at the contig-level reduced the number to 647 contigs. Structural annotation was performed on the contigs encoding 16S rRNA using Prodigal to generate protein sequences, which were subsequently used for the ML estimation. We then iterated a series of phylogenetic reconstructions by generating several supermatrix alignments, replacing the proteins from ptMAGs (n=1027 bins) instead of limiting to contigs with 16S rRNA genes, and adjusting the parameter of minimum numbers of PLASTID54 markers in the final alignment (genomes retaining at least 10%, 20%, 30%, and 40%). The ML species trees were then estimated for each supermatrix alignment using IQ-TREE with the LG+F+I+G4 substitution model, and branch support was evaluated with 1,000 ultrafast bootstrap pseudoreplications.

### Dereplicating and identifying novel ptMAGs

To eliminate redundant ptMAGs, we employed a similar approach as for the Cyanobacteriota, but with some modifications. First, we generated an initial ML tree for the ptMAGs, setting a minimum at 10% of PLASTID54 markers for inclusion in the final supermatrix alignment. Using the ML tree, we calculated pairwise evolutionary distances between cluster representatives with PhyloDM and at a 0.99 cutoff value with the inflation value of 1.5 using MCL clustering. Second, we generated cluster statistics and sorted the cluster based on the count of PLASTID54 markers. To represent each cluster, we selected plastid metagenomes based on the criteria of highest PLASTID54 markers count, fewest contigs per metagenomic bin, overall size of the contigs, and highest number of proteins.

To identify novel ptMAGs, we downloaded 15,248 complete plastid genomes available at NCBI RefSeq (https://ftp.ncbi.nlm.nih.gov/refseq/release/plastid/; last access October 2024) along with all plastid sequences ≥3,000 bp available in the NCBI GenBank (last access October 2024), creating a dataset of 28,983 publicly available plastid sequences, including complete and partial plastomes. We calculated average nucleotide identity (ANI) for the ptMAGs against the plastid dataset using skani with the “skani dist --slow” option. Plastid MAGs were classified as redundant based on two criteria: (i) ANI ≥95%, align_fraction_ref ≥ 50%, and align_fraction_query ≥ 50% (ptMAGs), and (ii) ANI ≥95% and align_fraction_query ≥95% with any values for the align_fraction_ref. The latter criterion specifically aimed to filter out smaller ptMAGs with very high ANI that only cover partial regions of a complete reference plastome.

### Annotation of plastid metagenomes

Protein-coding genes, ribosomal RNAs, and transfer RNAs in ptMAGs were annotated using GeSeq v2.03^55^, and tRNAscan-SE v2.07^56^, followed by manual curation using Geneious Prime 2025.0.2 (www.geneious.com). Whole-genome alignment was carried out using progressiveMauve^57^ within the Geneious platform.

## RESULTS AND DISCUSSION

### Expanding the cyanobacterial taxonomy for tracing plastid origins

To investigate the origin of plastids, we first expanded the phylogenetic framework of Cyanobacteriota taxonomy. With the extended MAG binning, a total of 1,973 genomes/MAGs belonging to Cyanobacteria (referred to as Cyanobacteriia in the GTDB), Vampirovibrionia, and Sericytochromatia were analyzed. This dataset included 438 new metagenomic bins along with 1,539 genomes/metagenomes available in two public databases: the GTDB and the (IMG/M) databases. Among the 438 new metagenomic bins, GTDB taxonomic classification assigned eight, 84, and 346 bins into Sericytochromatia, Vampirovibrionia, and Cyanobacteria, respectively (Supplementary Table S2). Two groups, Sericytochromatia and Vampirovibrionia, were exclusively represented by metagenomic data, while the Cyanobacteria included metagenomic bins as well as 496 genomes assembled from isolates (Supplementary Table S2). The inclusion of 438 new metagenomic bins within Cyanobacteriota increased the overall phylogenetic diversity by 28%. Our phylogenetic analyses strongly support the monophyly of the photosynthetic clade Cyanobacteria and the non-photosynthetic clades, Sericytochromatia and Vampirovibrionia, with Sericytochromatia positioned at the base of the clade and sister to Vampirovibrionia and Cyanobacteria (Fig. 1a, Supplementary Fig. S1). This topology is consistent with published studies^29,58^. For detailed description of the Cyanobacteriota phylogeny, see Supplementary Text.

**Figure 1.**
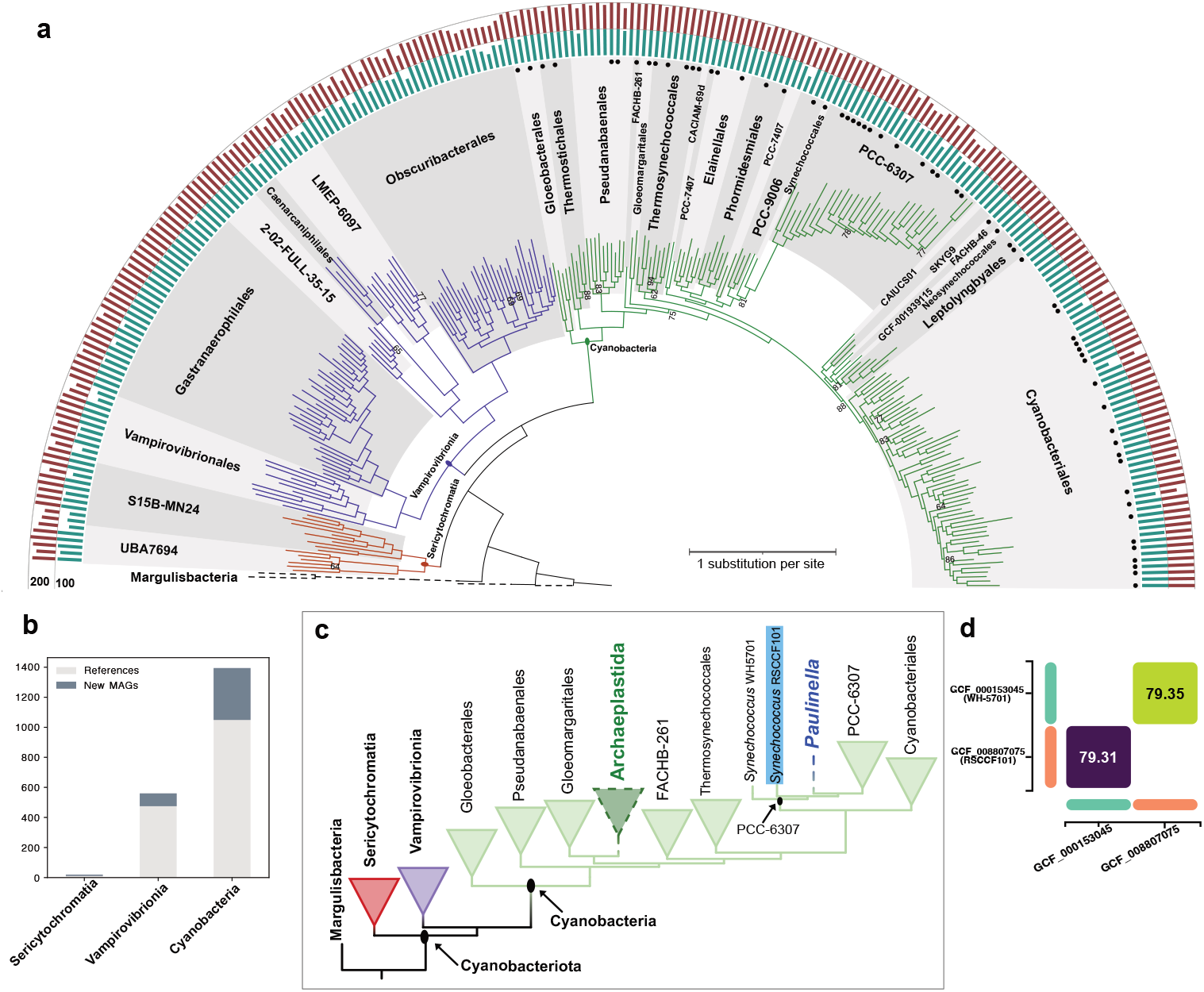
Phylogenic placement of primary plastids within the cyanobacterial lineage. (a) A maximum-likelihood tree of the dereplicated Cyanobacteriota dataset, constructed using concatenated UNI56 markers with IQ-TREE. Branches are color-coded: deep red for Sericytochromatia, violet for Vampirobivrionia, and green for Cyanobacteria. The tree is rooted using Margulisbacteria. Taxonomic orders within are labeled and distinguished with different shades of grey. Only ultrafast bootstrap values <95% are shown, either above or below the branches. Isolates are marked with filled black circles at the end of the leaves, distinguishing them from MAGs. The outside bar plots represent the count of UNI56 markers (dark cyan) and CheckM2 completeness (dark red). (b) A stacked histogram showing the count of publicly available and newly generated MAGs for Cyanobacteriota in this study. (c) A simplified illustration showing the phylogenetic positions of primary plastids (Archaeplastida and *Paulinella*) within the cyanobacterial phylogeny. Selected cyanobacterial taxonomic orders are shown as collapsed triangles to ease visualization of plastid origins, which are highlighted by dotted clade and branch lines. The *Synechococcus* species RSCCF101, which is phylogenetically closer to the *Paulinella* plastid, is highlighted with a blue background. (d) Whole-genome ANI comparison between *Synechococcus* WH-5701 and *Synechococcus* RSCCF101.

To determine the phylogenetic placement of plastids within the Cyanobacteriota taxonomy, we conducted a series of phylogenetic reconstructions by varying the datasets and the phylogenetic markers used for the supermatrix alignments (detailed description of the iterated analyses can be found in methods). In all phylogenetic reconstructions, regardless of increasing taxon sampling, swapping the phylogenetic markers, or modifying cut-off parameters for tree reconstructions, we observed that plastomes and ptMAGs grouped together in one of two strongly supported monophyletic clades (UFBoot 100%) within the Cyanobacterial tree (Fig. 1c, Supplementary Figs. S2). The first clade represents the primary plastids of *Paulinella* and confirms relatedness to an ancestor of the α-cyanobacterial clade containing *Synechococcus* species^11,12^. However, while *Synechococcus* sp. WH5701 was the assumed closest extant relative^12^, our results place the *Paulinella* plastid within the cyanobacterial order PCC-6307, after the diversification of *Synechococcus* sp. RSCF101 (Fig. 1c, Supplementary Fig. S3). The *Synechococcus* sp. RSCF101 shared ∼79% nucleotide identity with *Synechococcus* sp. WH5701, and the two *Synechococcus* species belonged to distinct phylogenetic clades within the PCC-6307 (Fig. 1d, Supplemental Fig. S3). The second clade contains all plastomes and ptMAGs other than *Paulinella* and is sister to the cyanobacterial order Gloeomargaritales (Fig. 1d, Supplementary Fig. S2). This finding supports the current understanding that primary plastids in Archaeplastida originate from a deeply branching cyanobacterium closely related to *Gloeomargarita* lineage^6,7,59^, and does not support any alternative scenarios^60,61^.

### An extended taxonomic framework of plastids tracing novel ptMAG lineages

Next, we investigated the relationships among plastids in photosynthetic eukaryotes and the impact of our novel ptMAGs on established plastid phylogenetic relationships. The resulting trees revealed a consistent topology of major photosynthetic eukaryotes within green and red lineages, such as Streptophyta, Chlorophyta, Rhodophyta, Cryptophyta, Haptophyta, and Ochrophyta. Within the green lineage, Prasinodermophyta emerged as an early diverging clade and sister to Chlorophyta and Streptophyta with varying UFBoot support values (93% to 100%), substantiating their classification as a new phylum within Viridiplantae^62,63^. Streptophytes and chlorophytes were strongly supported as monophyletic groups, and lineages with secondary/complex plastids, such as Euglenophyceae, Chlorarachniophytes, and the dinoflagellates species *Lepidodinium*, which acquired plastids from unrelated green algae, were included within chlorophytes. The two strongly supported clades of plastids from green lineages (Streptophyta and Chlorophyta) were recovered as a sister to the plastids in red lineage (Rhodophyta, Cryptophyta, Haptophyta, Ochrophyta) and Glaucophyta (UFBoot 100%; Fig. 2, Supplementary Figs. S4 & S5). Our results strongly supported Glaucophyta plastids as a sister group to the red lineages plastids (UFBoot >95%; Fig. 2, Supplementary Figs. S4 & S5). This relationship is not uncommon, and it is known that glaucophytes share more plastid-encoded genes with rhodophytes than with plastids from green lineage^64,65^. Ochrophyta also included kleptoplasty-derived complex plastids in two unrelated lineages: a marine centroplasthelida *Meringosphaera mediterranea* (a Haptista), and two dinoflagellate species, *Kryptoperidinium foliaceum* and *Durinskia baltica*.

**Figure 2.**
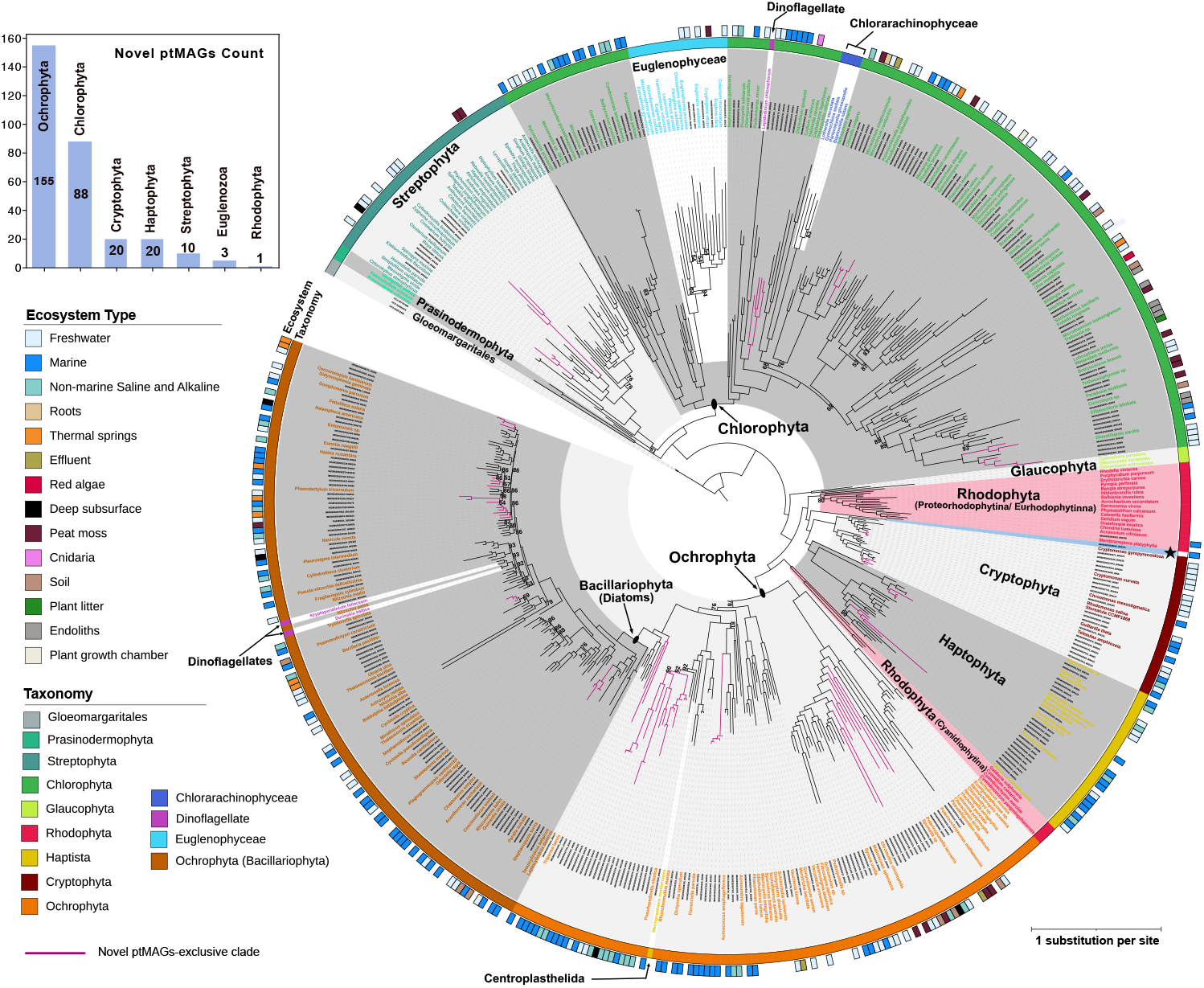
Plastid relationships among major photosynthetic eukaryotes. The maximum-likelihood tree was constructed using a concatenated alignment of PLASTID54 markers under the LG+F+I+G4 substitution model, following the nsgtree pipeline and IQ-TREE analysis. The dataset comprises 536 taxa, including 236 reference plastomes (the font of species names is colored according to the “Taxonomy” color key), 295 ptMAGs (labeled with black font), and three Gloeomargaritales species used as an outgroup to root the tree. The bar plot insert illustrates the number of novel ptMAGs identified per major photosynthetic group. The inner ring indicates the assigned taxonomy for each clade, as described in the color key. Different background shades of gray are used to distinguish separate clades; clades representing secondary/complex green algal plastids in Euglenophyceae, Chlorarachinophyceae, and the dinoflagellate *Lepidodinium chlorophorum*, and complex red algal plastids in dinoflagellates and Centroplasthelida are excluded from the shaded areas. The outer ring indicates the ecosystem of origin for the ptMAGs. A single ptMAG, which diverges before the diversification of Haptophyta and Cryptophyta, is highlighted with blue background shading and marked with a black star. Ultrafast bootstrap values are displayed only for branches with support below 95%. Some examples of clades represented exclusively by two or more ptMAGs are highlighted with magenta branches.

Interestingly, in all our phylogenetic reconstructions, the red algae lineage is split (Fig. 2, Supplementary Figs. S4 & S5). The mesophilic red algae were monophyletic and strongly supported as a sister to Cryptophyta and Haptophyta (Fig. 2). Whereas the early diverging red algal subphylum Cyanidiophytina was strongly supported as a sister to Ochrophyta (Fig. 2). This finding is consistent with the initial observation by Kim et al. (2015)^66^, and a more recent study by Pietluch et al. (2024)^67^. Their study, like ours, presented a nearly identical topology, suggesting two independent origins of secondary red algal plastids and proposing a “multiple and serial endosymbiosis” model. This model extends the serial endosymbiosis hypothesis, which posits an initial secondary endosymbiosis of red algae plastids by cryptophytes, followed by acquisition of cryptophytes plastids by ochrophytes and haptophytes following tertiary or higher order endosymbiosis^19–21^. Our study, along with the findings of Pietluch et al. (2024), challenges the notion of a single origin for secondary red algal plastids and the vertical inheritance of the plastids in all lineages with red algal-derived secondary/complex plastids, presenting evidence for two distinct engulfment events involving a red alga.

Our comprehensive phylogenomic analysis not only expands the known plastid diversity across most photosynthetic lineages but also reveals previously unidentified groups. We examined the phylogenetic distribution of 300 novel ptMAGs and compared them to selected RefSeq plastomes from various photosynthetic groups (Fig. 2). Since these novel ptMAGs were obtained using a dereplicated ptMAGs dataset, the true diversity of the novel ptMAGs is likely underrepresented. Among the novel ptMAGs, the largest number belonged to Ochrophyta (n=155), followed by Chlorophyta (n=88), Haptophyta (n=20), Cryptophyta (n=20), Streptophyta (n=10), Euglenophyceae (n=5) and Rhodophyta (n=1) (Supplementary Table S5, Fig. 2). Interestingly, we identified a novel ptMAG as a sister to Cryptophyta and Haptophyta (discussed in more detail below). We did not recover any ptMAGs representing Glaucophyta and Prasinodermaphyta, and both phyla were solely represented by RefSeq plastomes.

The ptMAGs identified in our study reveal previously unreported plastid diversity, highlighting several noteworthy examples that likely represent undescribed algal lineages. Within the green lineages, assembly sizes for Streptophyta ptMAGs ranged from ∼61 Kb to ∼131 Kb, and ptMAGs (n=6) recovered from freshwater and deep subsurface formed a monophyletic clade sister to charophyte algae, suggesting they are new members of class Zygnematophyceae (Fig. 2). In the Chlorophyta, the novel ptMAGs numbers were substantially larger, with sizes ranging from ∼20 Kb to ∼209 Kb. Among these, potential novel lineages include ptMAGs (n=5) from marine ecosystems forming a monophyletic clade sister to Pedinophyceae, and ptMAGs (n=4) from marine and freshwater forming a distinct clade within Trebouxiophyceae. A single novel rhodophyte ptMAG of size ∼171 Kb was recovered as a sister to *Membranoptera platyphylla* (family Delesseriaceae) plastome, and given the rarity of red algae plastomes, the novel ptMAG will be valuable. Additionally, within Cryptophyta and Haptophyta, 20 novel ptMAGs were identified in each group, with sizes ranging from ∼96 Kb to ∼150 Kb and from ∼10 Kb to ∼138 Kb, respectively. Within Cryptophyta, novel ptMAGs (n=9) from freshwater habitats grouped with RefSeq *Cryptomonas* species, while a clade with six ptMAGs (n=6) from diverse habitats were placed in the clade with Geminigeraceae species. In Haptophyta, novel ptMAGs from marine, freshwater, and non-marine saline and alkaline habitats were placed in a clade with *Pavlova* species (ptMAGs n=6), and another clade containing Chrysochromulinaceae plastids (ptMAGs n=9). Both Cryptophyta and Haptophyta included a limited number of complete plastomes in public databases, and the novel ptMAGs provide essential plastid diversity for these clades.

Given that many novel ptMAGs identified were from Ochrophyta, we performed an extensive phylogenomic analysis to determine their distribution within the phylum, with detailed findings presented in Supplementary Text. Here, we briefly summarized some notable results. We identified a total of 155 novel ptMAGs within Ochrophyta, with sizes ranging from ∼19 Kb to ∼141 Kb (Supplementary Table S5, Supplemental Figs. S6 & S7). The majority of these ptMAGs (n=122), originating from diverse habitats including freshwater, marine, deep subsurface, non-marine saline and alkaline, thermal springs, and soil, were classified under Bacillariophyta, showcasing significant plastid diversity associated to diatoms. In addition, we identified four novel ptMAGs of > 100 Kb within Bolidophyceae, a class sister to Bacillariophyta and represented by a single reference plastome in the public database^68^. Notably, we found a monophyletic clade consisting of three ptMAGs from marine habitats and non-marine saline and alkaline habitats, sister to Bacillariophyta and Bolidophyceae. Despite their smaller size (<50 Kb), these ptMAGs could be key to advancing our understanding of plastid evolution in diatoms. Other classes with a substantial number of ptMAGs identified include Synurophyceae (n=27), Dictyochophyceae (n=24), and Pelagophyceae (n=10). Many ptMAGs within these classes are from diverse habitats, such as freshwater, marine, deep subsurface, soil, peat moss, non-marine saline and alkaline, and effluent, and formed clades without reference plastomes.

The newly recovered ptMAGs from this study, especially those within Chlorophyta and Ochrophyta, hold considerable promise to clarify plastid relationships and evolution across different taxonomic levels. To accomplish this, it will be necessary to use lineage-specific markers by carefully selecting informative genes to improve phylogenetic resolution. Additionally, employing suitable evolutionary models and phylogenetic techniques will be crucial for tackling difficult lineages.

### New lineage sister to Cryptophyta and Haptophyta

We identified a novel ptMAG from a marine ecosystem (Artic Ocean near Svalbard) that exhibits an intriguing phylogenetic relationship with Cryptophyta and Haptophyta. This ptMAG (IMGM3300009544_BIN141) is sister to mesophilic red algae and branches before the last common plastid ancestor of Cryptophyta and Haptophyta. This ptMAG could potentially provide new insight into the acquisition of plastids from red algae in Cryptophyta before the spread to Haptophyta. The ptMAG consists of a single 104,344 bp contig likely representing the large single copy region of the plastome with partial 23S rRNA and 16S rRNA gene sequences at either end and retains 108 protein-coding genes with 26 tRNAs genes (Supplemental Table S6). The identified genes include those encoding large and small ribosomal subunits (38 genes), photosystem I/II reaction center (25 genes), ATP synthase (8 genes), cytochrome *b*_6_*f* complex (6 genes), plastid-encoded RNA polymerase (4 genes), carbon fixation (3 genes), cytochrome *c* biogenesis (2 genes), and protein translocase (2 genes). The ptMAG lacks several plastid genes found in haptophytes and/or cryptophytes, such as *ftsH, minE, dnaB/X*, and *chlN/L/B*^69^, but we cannot rule out that the absences are due to the partial nature of the assembly. However, the ptMAG contains two genes, *minD* (encoding septum site-determining protein) and eubacterial *c*-type *rpl36* (encoding the large ribosomal subunit protein L36, with the best BLAST hit to RPL36 protein from bacterial order Pirellulales), which are absent in the plastomes of mesophilic red algae but shared by plastomes of cryptophytes and haptophytes. The presence of eubacterial *c*-type *rpl36* in cryptophytes and haptophytes plastomes is attributed to horizontal gene transfer in an ancestor of cryptophytes before engulfment of a cryptophyte by the haptophyte ancestor^70^. This ptMAG could come from an algal lineage that diverged prior to acquisition of the plastid by haptophytes, or this ptMAG is from a red alga that unlike other red algae encodes a Pirellulales *rpl36*.

Since our study includes only a single sequence of this novel ptMAG, we searched public databases for other sequences sharing high similarity. Initially, we used the partial 16S rRNA gene sequence from the ptMAG to search for similar sequences in the NCBI database. The nucleotide BLAST search revealed that the closest match is a 16S rRNA gene sequence (KX935025; ∼92% nucleotide pairwise identity) from an uncultured marine eukaryote in the DPL2 clade identified by Choi et al. (2017)^71^. DPL2 (Deep-branching Plastid Lineage 2) is a globally distributed deep-branching clade positioned as a sister to haptophytes^71^. We also used the protein sequences for plastid genes from the ptMAG IMGM3300009544_BIN141 to search the MGnfiy microbiome database^63^ (https://www.ebi.ac.uk/metagenomics) with phmmer. We identified several assemblies with significant e-values (1 × 10^−208^ or lower), mainly corresponding to Tara Oceans assemblies and a few from a marine microbial diversity study from Saanich Inlet. Several contigs were identified with sizes ranging from ∼1 Kb to ∼104 Kb and sharing >99% nucleotide identity with the novel ptMAG, however, none were longer than our assembly. The longest contigs from that Tara Oceans assemblies (TARA_B110000971 and TARA_B110000977, from different Artic Ocean locations) shared identical gene content with our novel ptMAG IMGM3300009544_BIN141, including 100% amino acid identity with eubacterial *c*-type RPL36. The 16S rRNA gene sequence from all these sequences exhibit the highest sequence similarity (>90%) with DLPL2 16S rRNA sequence. The highly similar genomic features between the Tara Oceans contigs and our novel ptMAGs suggest that they are likely all from closely related, undescribed marine algae.

Our findings were further corroborated by a recent preprint by Jamy et al. (2025)^72^, which describes a new algal group based on plastid sequences referred to as “Leptophytes” and places them deep within the algal tree of life. Our novel ptMAG IMGM3300009544_BIN141 shares 99.9% identity with the best representative plastome for Leptophytes (Lepto-01)^72^. The phylogenetic placement of the novel ptMAG as a sister to both cryptophytes and haptophytes was supported by Jamy et al. (2025). However, an alternative topology where leptophyte plastids closer to haptophyte plastids than cryptophytes was also presented by authors creating uncertainty about their exact phylogenetic placement. Much like our ongoing efforts, the authors were unable to identify the host of leptophytes. Nonetheless, the novel ptMAG offers an essential plastid sequence that could potentially bridge the evolutionary gap between red algae and Cryptophyta/Haptophyta.

### Novel ptMAGs linked to secondary/complex green-algal plastids

Complex plastids derived from green algae have been previously identified in three distinct lineages: Chlorarachniophyceae, Euglenophyceae, and the dinoflagellate species *Lepidodinium chlorophorum*. Notably, in *L. chlorophorum*, the original plastid is believed to have been replaced by a chlorophyte-derived plastid^15,73–75^. Our phylogenetic analyses support the independent origins of green alga-derived secondary plastids in these three lineages (Fig. 2, Supplementary Figs. S4 & S5). No novel ptMAGs were found within Chlorarachniophyceae; this clade was solely represented by the four RefSeq plastomes of *Lotharella vacuolata, Gymnocholora stellata, Bigelowiella natans*, and *Partenskyella glossopodia*. Chlorarachniophyceae formed a strongly supported monophyletic group, sister to the order Trentepohliales, (moderate to high support values; UFBoot 77% to 98%). Chlorarachniophyceae together with Trentepohliales was strongly supported as sister to order Bryopsidales, in agreement with a previous study that concluded siphonous green algae are likely ancestors of Chlorarachniophyceae plastids^15^.

Our study consistently recovered secondary plastids in Euglenophyceae as a sister clade to the prasinophyte algae order Pyramimonadales with strong support. In some cases, euglenophyte plastids were placed closer to *Cymbomonas tetramitiformis* than to other Pyramimonadales, albeit with moderate support (UFBoot 85%; Fig. 2, Supplementary Figs. S4 & S5). Although the specific Pyramimonadales alga closest to euglenophytes has not been determined, the sister relationship between Pyramimonadales and euglenophytes has been consistently recovered in previous studies. Our findings further support that secondary plastids in euglenophytes share a common ancestor with Pyramimonadales algae^65,74,76^. We identified several novel ptMAGs both within Pyramimonadales and Euglenophyceae. Specifically, novel ptMAGs (n=5) with assembly sizes ranging from ∼44 Kb to ∼104 Kb were discovered within Euglenophyceae, and ptMAGs (n=4) sizes ranging from ∼70 Kb to ∼90 Kb were placed within Pyramimonadales.

Among the lineages with chlorophyte-derived secondary plastids, we uncovered several novel ptMAGs closely related to *L. chlorophorum* (Fig. 2). To gain insights into the origin of plastids in this dinoflagellate, we conducted comparative analyses (Fig. 3). The plastid origin in the *L. chlorophorum* has been attributed to an endosymbiotic event involving a pedinophyte algae closely related to *Pedinomonas minor*^15,77^. We identified two novel ptMAGs more closely related to *L. chlorophorum* than *Pedinomonas* species. To determine the exact position of these novel MAGs, we included additional Pedinophyceae plastomes available in the GenBank and reconstructed a phylogeny (Fig. 3a). Two novel ptMAGs, IMGM3300043446_BIN543, and IMGM3300027621_BIN154, with assembly sizes of ∼63 Kb and ∼90 Kb, respectively, were strongly supported as sister to *L. chlorophorum*. The ptMAG IMGM3300027621_BIN154 was represented by two contigs (∼81 Kb and ∼9 Kb), with the larger contig containing ribosomal rRNA operons at both ends, and the shorter contig containing genes, *cysT, ycf1, ccsA, psaC*, which are typically located within the small single copy of *Pedinomonas* species. The ptMAG IMGM3300043446_BIN543 consists of a single contig of ∼64 Kb and shares 65% ANI with the ptMAG IMGM3300027621_BIN154. The larger ptMAG IMGM3300027621_BIN154 likely represents a near complete assembly and was selected for the gene content analysis. It retained almost all protein-coding genes present in *L. chlorophorum* except for *petL* and *rpoA*, while also including nine additional genes (Fig. 3b). *L. chlorophorum*, along with the ptMAGs IMGM3300043446_BIN543 and IMGM3300027621_BIN154, was strongly supported as a sister to a clade containing two *Pedinomonas* plastomes and a novel ptMAG (IMGM3300042988_BIN337) of size 87,241 bp. The gene content of ptMAG IMGM3300042988_BIN337 was identical to *P. minor*, but three syntenic blocks were inverted at the location 64 Kb - 72 Kb (Fig. 3b-c). Considering the overall gene content, synteny, and the phylogenetic position, the ptMAG IMGM3300042988_BIN337 likely represents the plastome of a new *Pedinomonas* species. The *Pedinomonas* clade (*Pedinomonas* spp. and IMGM3300042988_BIN337) retained all the protein-coding genes present in *L. chlorophorum* and ptMAG IMGM3300027621_BIN154, along with six additional genes (Fig. 3b). Gene order in *L. chlorophorum* and its sister ptMAGs (IMGM3300043446_BIN543 and IMGM3300027621_BIN154) showed substantial rearrangements compared to *P. minor* (Fig. 3c). The ptMAGs IMGM3300027621_BIN154 and IMGM3300043446_BIN543 had similar gene order with inversions of syntenic blocks at three locations (4-12 Kb, 24-32 Kb, and 48-68 Kb). In contrast, the *L. chlorophorum* plastome was highly reduced and rearranged and gene order is not conserved. Additionally, the branch leading to *L. chlorophorum* was substantially longer compared to other plastomes and ptMAGs within Pedinophyceae. Considering shared gene content, synteny, and branch lengths, the ptMAGs IMGM3300027621_BIN154 and IMGM3300043446_BIN543 could potentially represent novel pedinophyte algae sister to *Pedinomonas*. If so, these two ptMAGs are more closely related to the *L. chlorophorum* plastid than the plastid from *P. minor*. It is also possible that these ptMAGs belong to novel dinoflagellate species that are sister to *L. chlorophorum* and have retained ancestral plastome features subsequently lost in *L. chlorophorum*.

**Figure 3.**
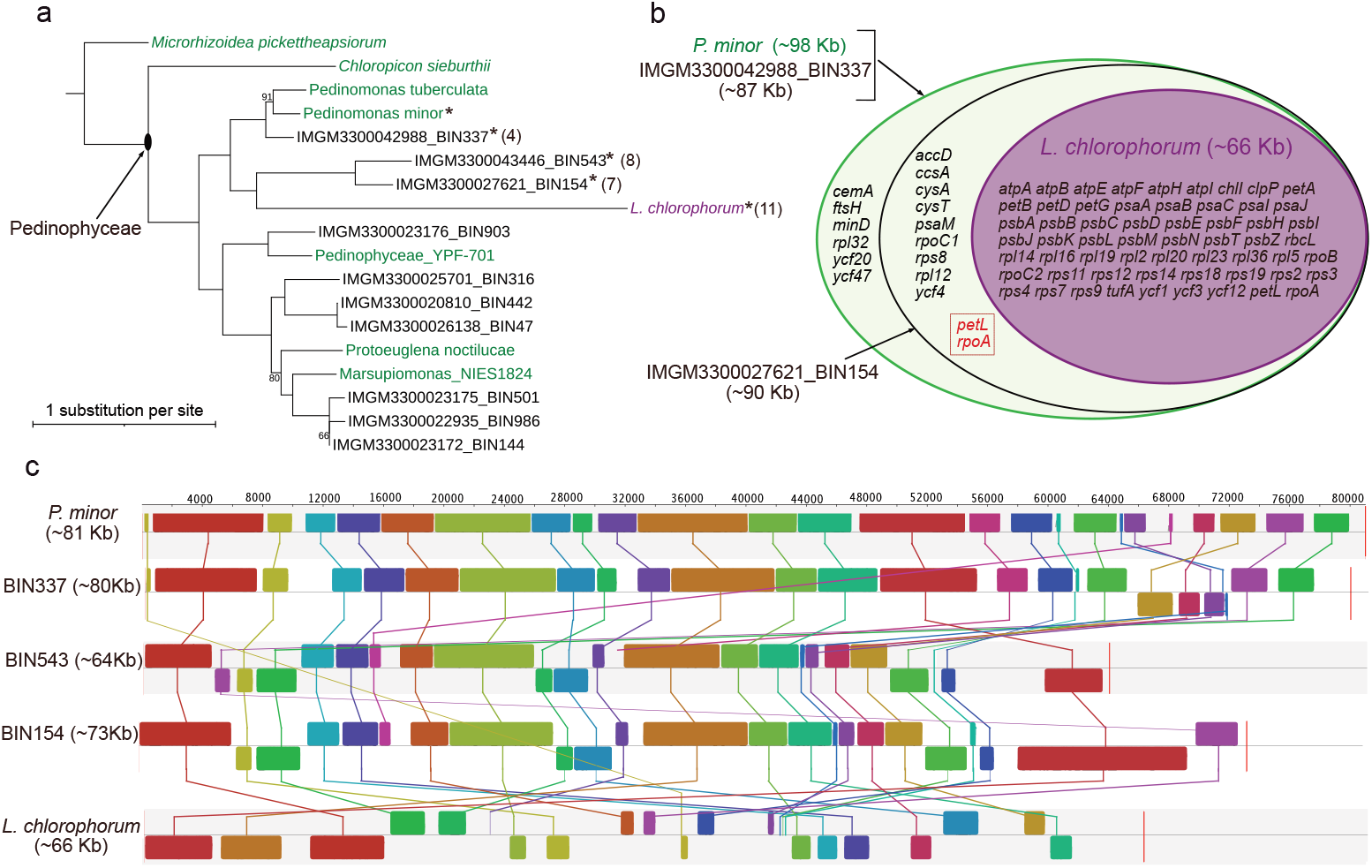
Closest relatives of green-algal-derived plastids in the dinoflagellate *Lepidodinium chlorophorum*. (a). Phylogenetic position of the *L. chlorophorum* plastid compared to *Pedinomonas* spp. plastids and novel ptMAGs. Additional Pedinophyceae plastomes available in GenBank are included. *Microrhizoidea* and *Chloropicon* spp. were used as an outgroup. Ultrafast bootstrap values <95% are shown. The reference species are color-coded, green for chlorophytes and purple for *L. chlorophorum*. Asterisks denote plastomes/MAGs used for whole genome alignment analysis. The number in the parentheses indicates the number of inverted syntenic blocks compared against *P. minor* (Fig. 3c). (b) A Venn diagram showing gene contents among *P. minor*, ptMAGs (IMGM3300042988_BIN337 and IMGM3300027621_BIN154), and *L. chlorophorum*, along with their respective plastome sizes. Two genes, *petL* and *rpoA*, missing only in IMGM3300027621_BIN154, are highlighted with a red dotted box. (c) Whole plastome alignment of ptMAGs and *L. chlorophorum* relative to *P. minor* generated using progressiveMauve alignment. Each locally collinear block representing synteny is color-coded; the blocks below the horizontal center indicate inversions. For simplicity, the ptMAGs labels have been abbreviated to associated bins. A copy of inverted repeats and single copy regions were removed from the plastome of *Pedinomonas* for the alignment.

## CONCLUSIONS

The endosymbiotic origin of primary plastids derived from cyanobacteria and their subsequent distribution among eukaryotes have led to remarkable diversity in photosynthetic eukaryotes. In this study, we explored the evolutionary relationships of primary plastids with cyanobacterial lineages and secondary/complex plastids among major algal groups by analyzing plastid sequences derived from metagenomic assemblies. By expanding the cyanobacterial taxonomic framework to include both photosynthetic and non-photosynthetic sister lineages, we pinpointed the origin of plastids. Our findings confirm that Archaeplastida plastids are closely related to the cyanobacterial order Gloeomargaritales, supporting the hypothesis that primary plastids originated from a deeply branching cyanobacterium, while *Paulinella* plastids arose independently. However, our results challenge existing models of plastid evolution in eukaryotes with secondary/complex plastids, presenting evidence for two independent origins of secondary red algae plastids. Recent studies align with this finding, emphasizing the need for further research and reassessment of red algal secondary endosymbiosis. Additionally, we uncovered numerous novel ptMAGs, particularly within Chlorophyta and Ochrophyta, revealing previously unexplored plastid diversity. Notably, we discovered a novel ptMAG that may represent an evolutionary link between red algae and Cryptophyta/Haptophyta, potentially tracing the common ancestor of secondary red algae plastids. The novel ptMAGs offers valuable insights into plastid evolution across photosynthetic eukaryotes.

## ACKNOWLEDGEMENTS

The work conducted by the U.S. Department of Energy Joint Genome Institute (https://ror.org/04xm1d337), a DOE Office of Science User Facility, is supported by the Office of Science of the U.S. Department of Energy operated under Contract No. DE-AC02-05CH11231. The work on chlorophyte algal plastids by B.S. and C.E.B-H. were supported by the Department of Energy (DOE) Office of Science, Biological and Environmental Research program under award no. DE-SC0023027. The work at the Molecular Foundry was supported by the Office of Science, Office of Basic Energy Sciences, of the U.S. Department of Energy under Contract No. DE-AC02-05CH11231.

## AUTHORS CONTRIBUTION

F.S. conceived the project. M.F.R. and J.C.V. generated MAGs associated with Cyanobacteriota and plastids and conducted data analysis. B.S. performed phylogenetic analysis and drafted the initial manuscript. C.E.B-H. and F.S. revised the manuscript. All authors read and approved the final manuscript.

## PLASTID MAG CONSORTIUM

Katherine McMahon, University of Wisconsin-Madison; Ramunas Stepanauskas, Bigelow Laboratory for Ocean Sciences; Alison Buchan, University of Tennessee-Knoxville; Thomas Mock, University of East Anglia; Kirsten Fisher, California State University; Joan Slonczewski, Kenyon College; and Luce Ward, Smith College.

## Notes

### Competing Interest Statement

The authors have declared no competing interest.

