## Supplementary Fig. S1 for "Global metagenomics reveals plastid diversity and unexplored algal lineages"

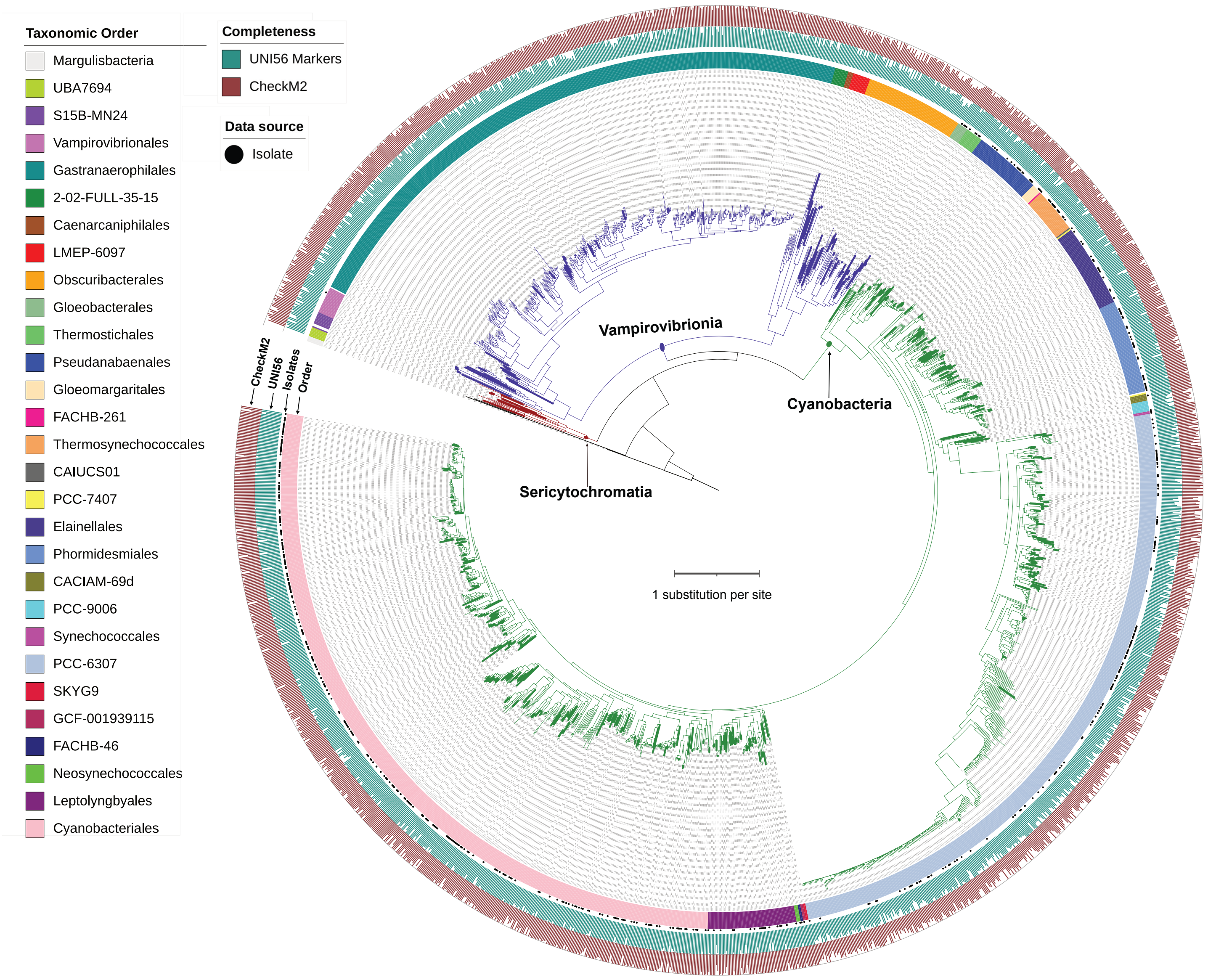


**Figure S1**. The phylogenetic relationships of 1,973 Cyanobacteriota genomes/MAGs were estimated using IQ-TREE, with Margulisbacteria as outgroup to root the tree. Each major clade is separated by color-coded branches: Sericytochromatia (dark red), Vampirovibrionia (dark blue), and Cyanobacteria (green). MAGs identified in this study (labeled as NeLLi2023 in Supplemental Table S2) are highlighted with thicker branches. Bootstrap support values are not shown. The diagram includes four rings: ring 1 indicates the GTDB taxonomic order for each cyanobacterial lineage; ring 2 uses filled circles to indicate isolates; ring 3 provides a bar plot showing UNI56 markers count; ring 4 provides a bar plot displaying CheckM2 completeness.
