## Supplementary Fig. S2 for "Global metagenomics reveals plastid diversity and unexplored algal lineages"

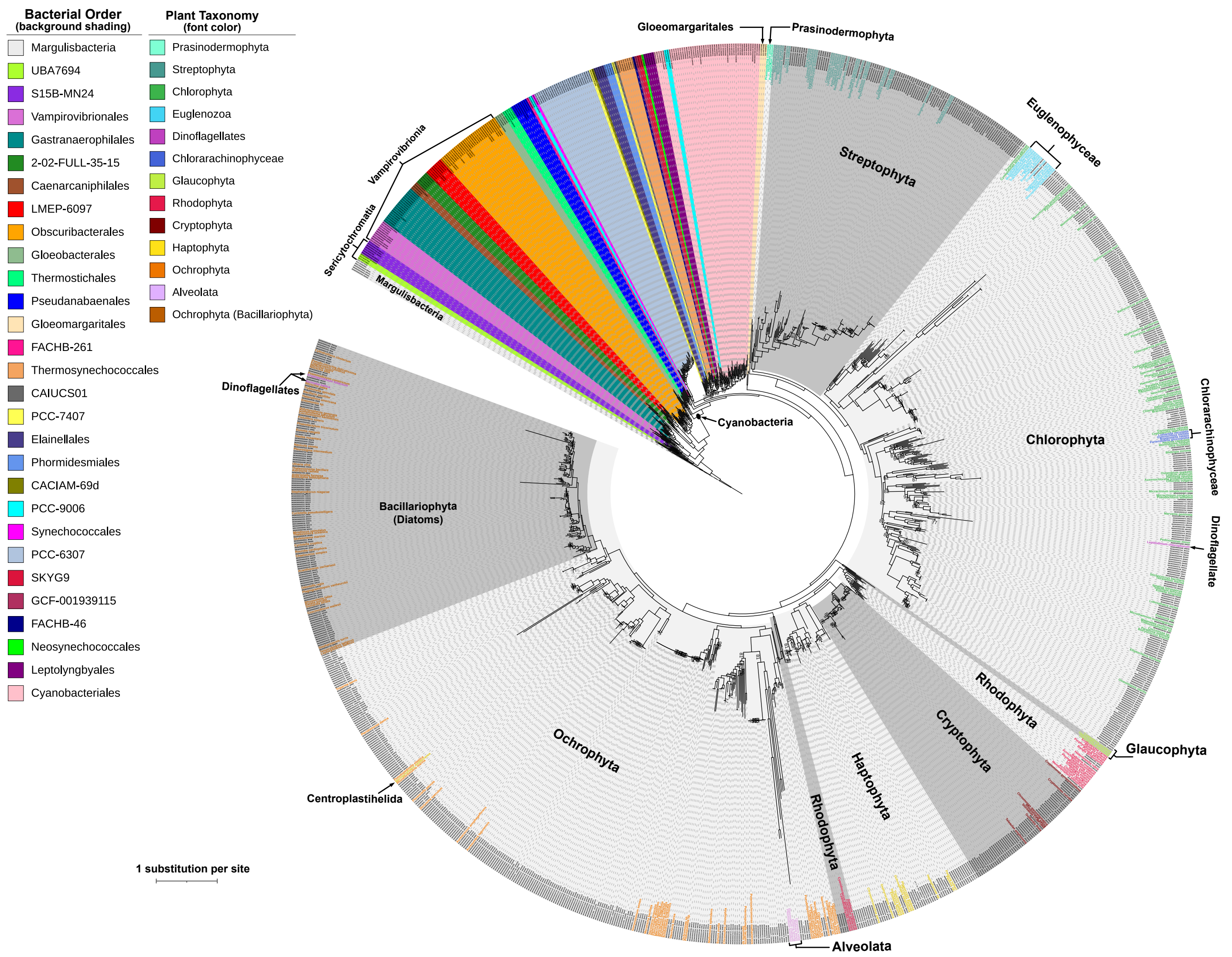


**Figure S2.** The phylogenetic relationships between cyanobacterial species and plastids from photosynthetic eukaryotes. Cyanobacterial species are represented by dereplicated 259 genomes/MAGs, while plastids are represented by 241 reference plastomes from major photosynthetic eukaryotic groups, and 647 ptMAG contigs containing 16S rRNA genes with at least 10% UNI56 markers. The tree was generated using a concatenated alignment of PLASTID54 markers under the LG+F + I + G4 substitution model with IQ-TREE. Cyanobacterial orders are highlighted with background color, with corresponding color codes for the clade provided in the color key (Bacterial Order). Gloeomargaritales, which is positioned as a sister to all plastids (reference plastomes and ptMAGs), is highlighted with an arrow and labeled. Labels for reference plastomes from diverse photosynthetic eukaryotes are color-coded and indicated in the color key (Plastid Taxonomy). Plastid metagenomes are indicated by labels with black text. Plastids from the major photosynthetic groups are separated with shades of gray and labeled by corresponding species name. The Alveolata clade includes *Chromera* species and four plastomes from Apicomplexa species but excludes dinoflagellates. Dinoflagellate members are separated from Alveolata to indicate their distinct phylogenetic positions.
