## Supplementary Fig. S3 for "Global metagenomics reveals plastid diversity and unexplored algal lineages"

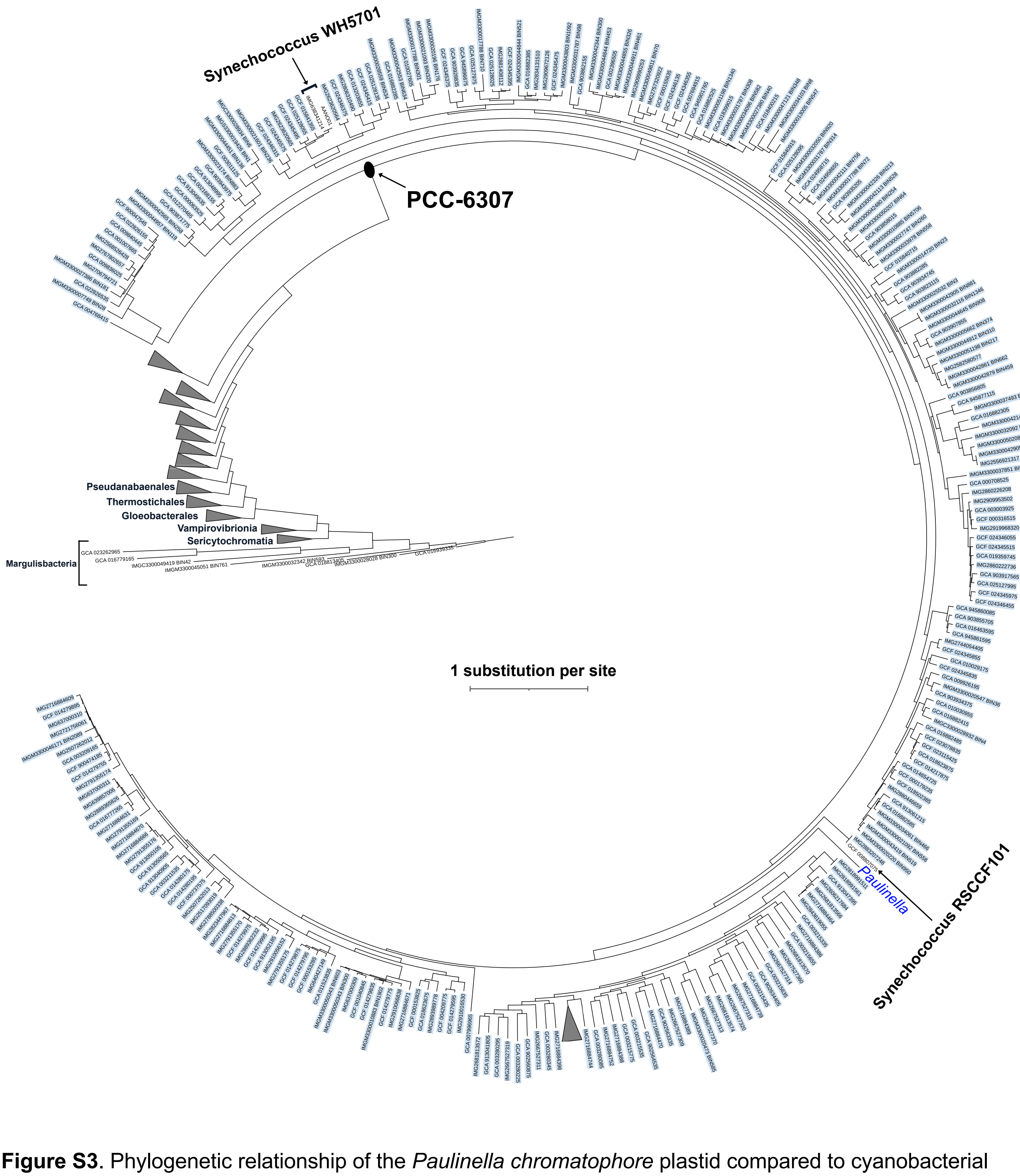


**Figure S3**. Phylogenetic relationship of the *Paulinella chromatophore* plastid compared to cyanobacterial members of the taxonomic order PCC-6307. The tree highlights the relationships of *Synechococcus* sp. WH5701 and *Synechococcus* sp. RSCCF101 in comparison to the *P. chromatophore* plastid. *Synechococcus* RSCCF101 branches off before the diversification of *P. chromatophore* plastid and is phylogenetically closer to *P. chromatophore*. Only members within the taxonomic order PCC-6307 are shown, while other orders within Cyanobacteriota are collapsed. For a complete overview of the relationship among all orders within Cyanobacteriota, please refer to Fig. 1 and Supplementary Fig. S1.
