## Supplementary Fig. S4 for "Global metagenomics reveals plastid diversity and unexplored algal lineages"

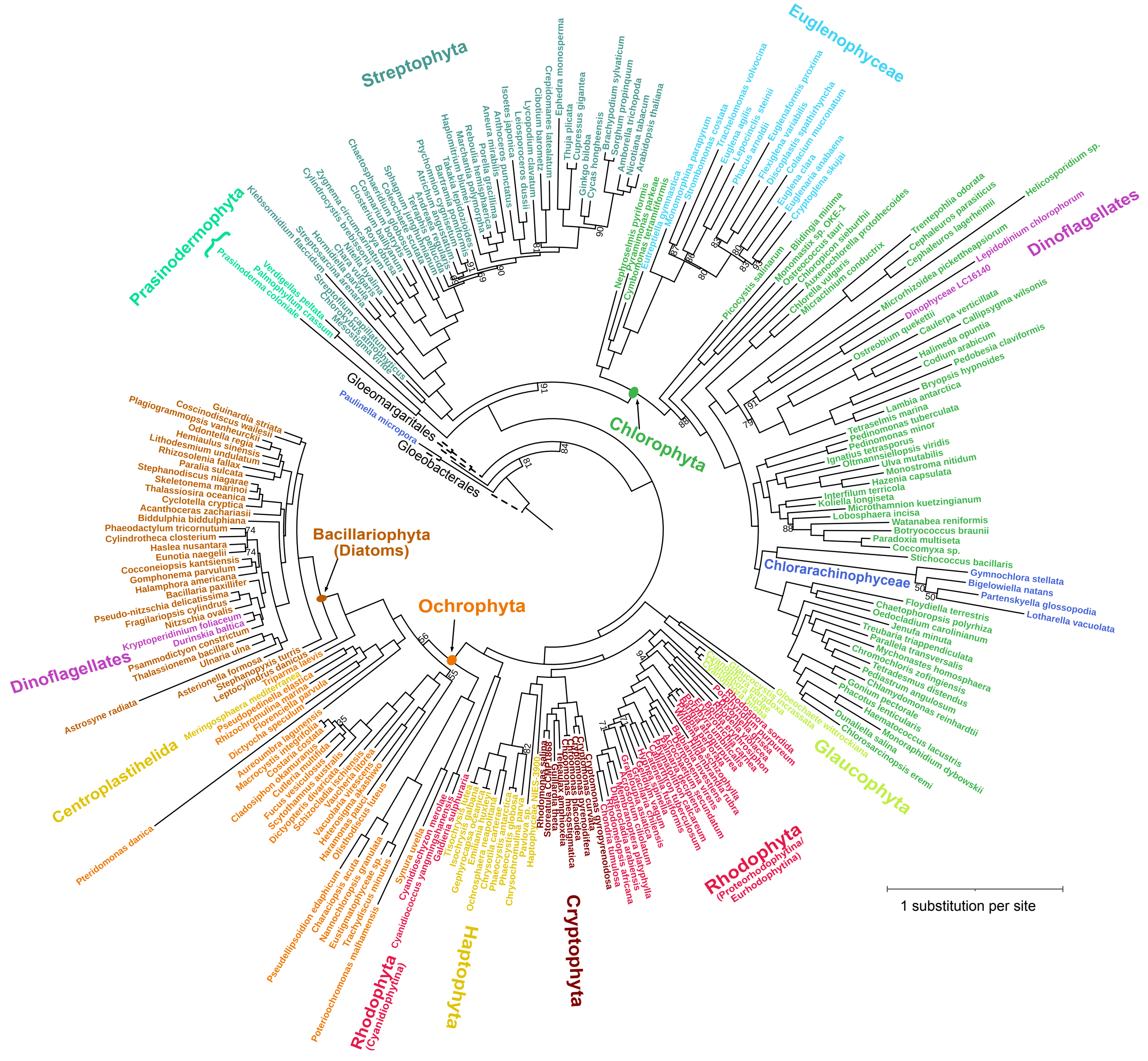


**Figure S4**. Phylogenetic relationships among different photosynthetic eukaryote plastids estimated with concatenated PLASTID54 genes using IQ-TREE. The dataset includes 236 RefSeq plastomes, *Paulinella chromatophore* plastid, and six cyanobacterial genomes. Ultrafast bootstrap values are displayed only for branches with support below 95%. Leaf labels are colored according to the color key.
