## Supplementary Fig. S5 for "Global metagenomics reveals plastid diversity and unexplored algal lineages"

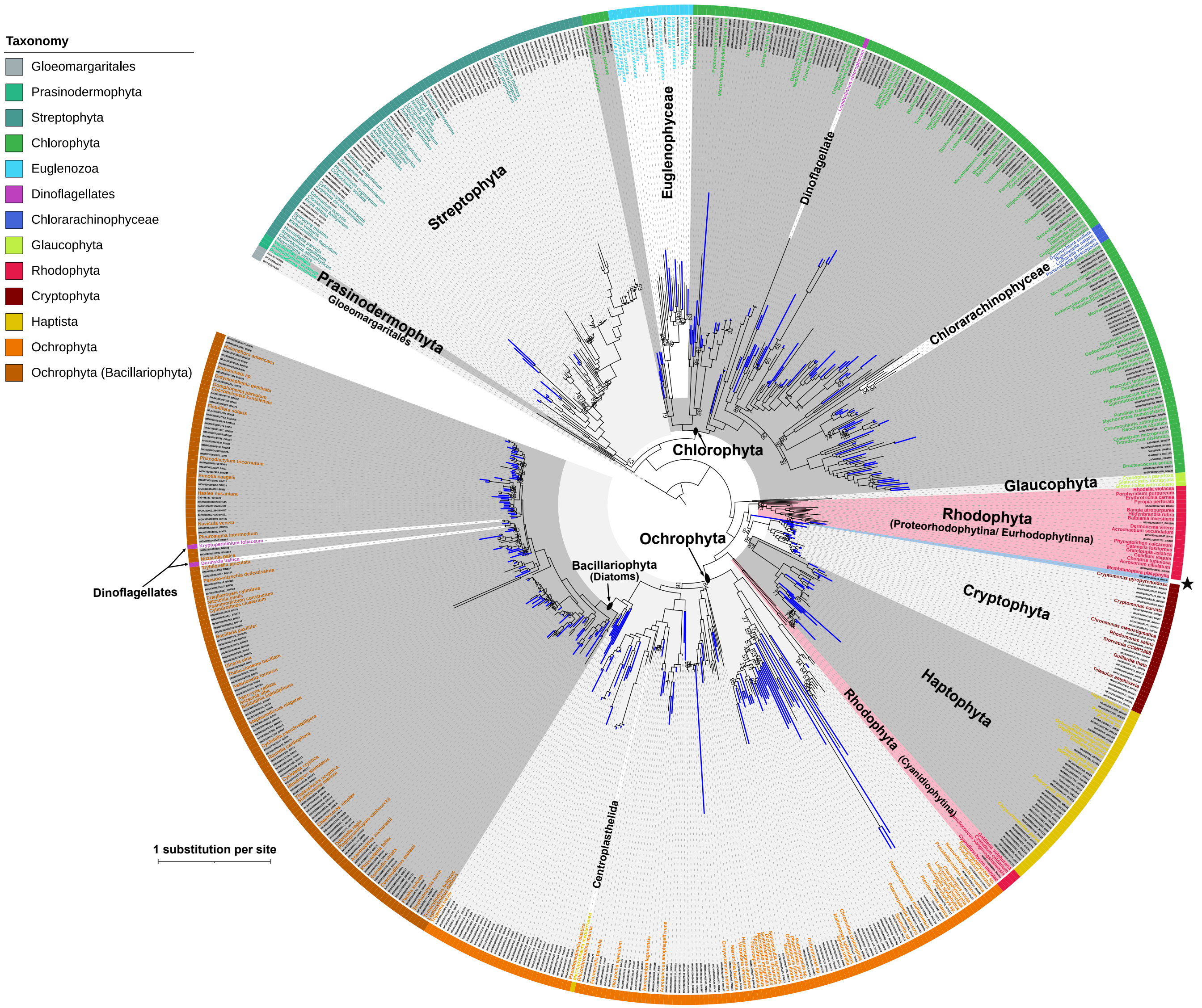


**Figure S5.** Plastid relationship among different photosynthetic eukaryotes. The maximum-likelihood tree was inferred using a concatenated alignment of PLASTID54 markers under the LG+F+I+G4 substitution model, following the nsgtree pipeline and IQ-TREE. Labels for the 236 reference plastids are color-coded as described in the color key, and labeled clades are delineated from one another with different shades of gray. Complex plastids within Chlorophyta (Euglenophyceae, Chlorarachinophyceae, and *Lepidodinum* spp.), and in dinoflagellates and Centroplasthelida are excluded from the shades. Rhodophyta clades are shaded in red to highlight two independent origins of secondary red algal plastids. A single plastid metagenome that branched off before the diversification of Haptophyta and Cryptophyta is highlighted in a blue shade and indicated by a black star. Novel ptMAGs are highlighted with bright blue branches to distinguish them from redundant ptMAGs derived in this study. Ultrafast bootstrap values below 95% are shown under the branches. Reference plastids from the phylum Bacillariophyta are shown in dark orange to distinguish from the rest of other ochrophytes (orange).
