## Supplementary Fig. S6 for "Global metagenomics reveals plastid diversity and unexplored algal lineages"

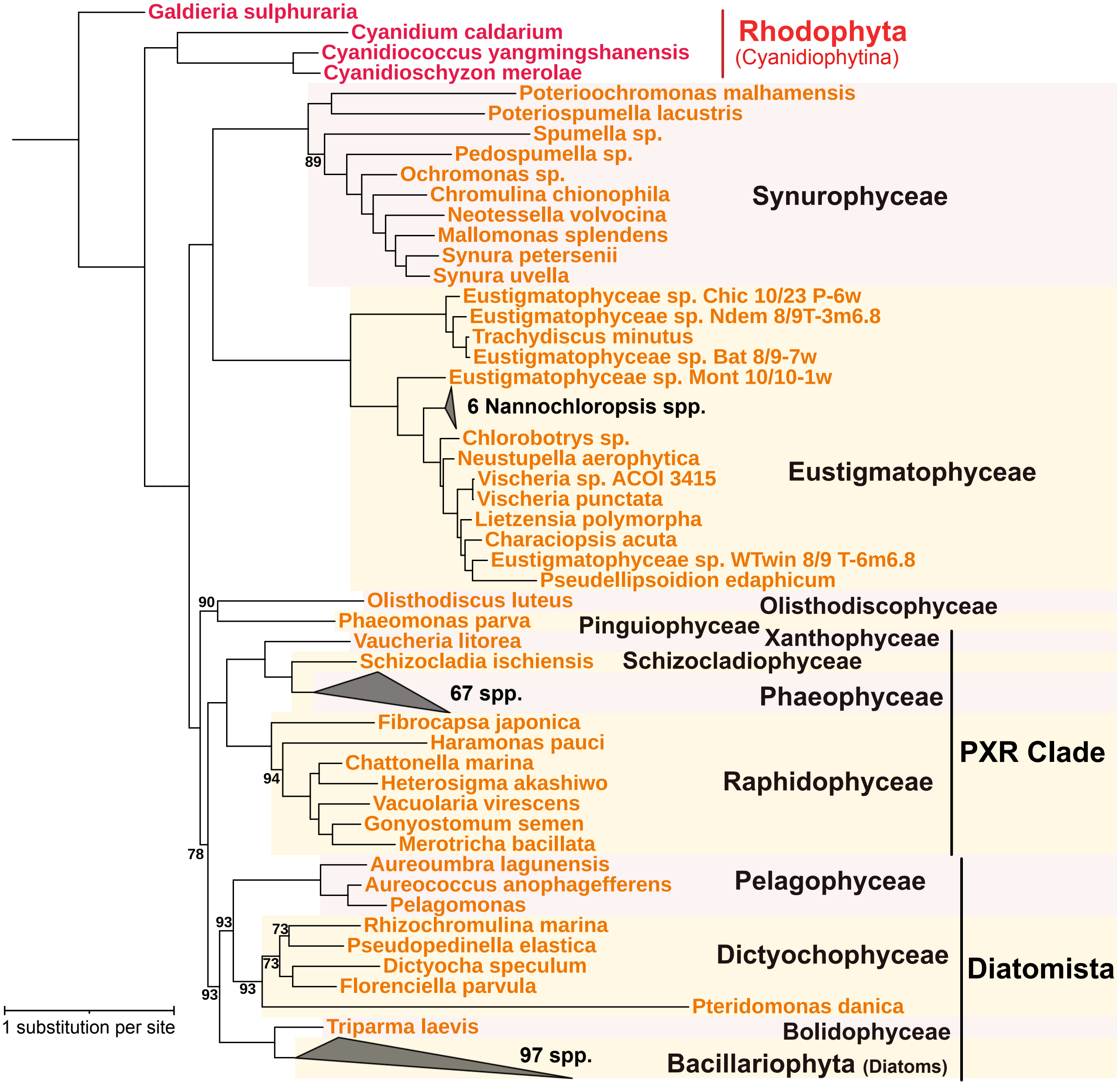


**Figure S6.** Phylogenetic relationships among reference Ochrophyta plastids. A maximum-likelihood tree was constructed with a concatenated alignment of 54 plastid proteins, applying the LG+F+I+G4 substitution model. The analysis included a total of 213 Ochrophyta reference plastids, with four Rhodophyta plastids serving as outgroup. Only ultrafast bootstrap support values less than 95% are displayed, either above or below the corresponding branches. Clades containing a large number of taxa were collapsed and labeled accordingly. Major groups within Ochrophyta were distinguished with alternating pink or yellow shading and labeled accordingly.
