## Supplementary Fig. S7 for "Global metagenomics reveals plastid diversity and unexplored algal lineages"

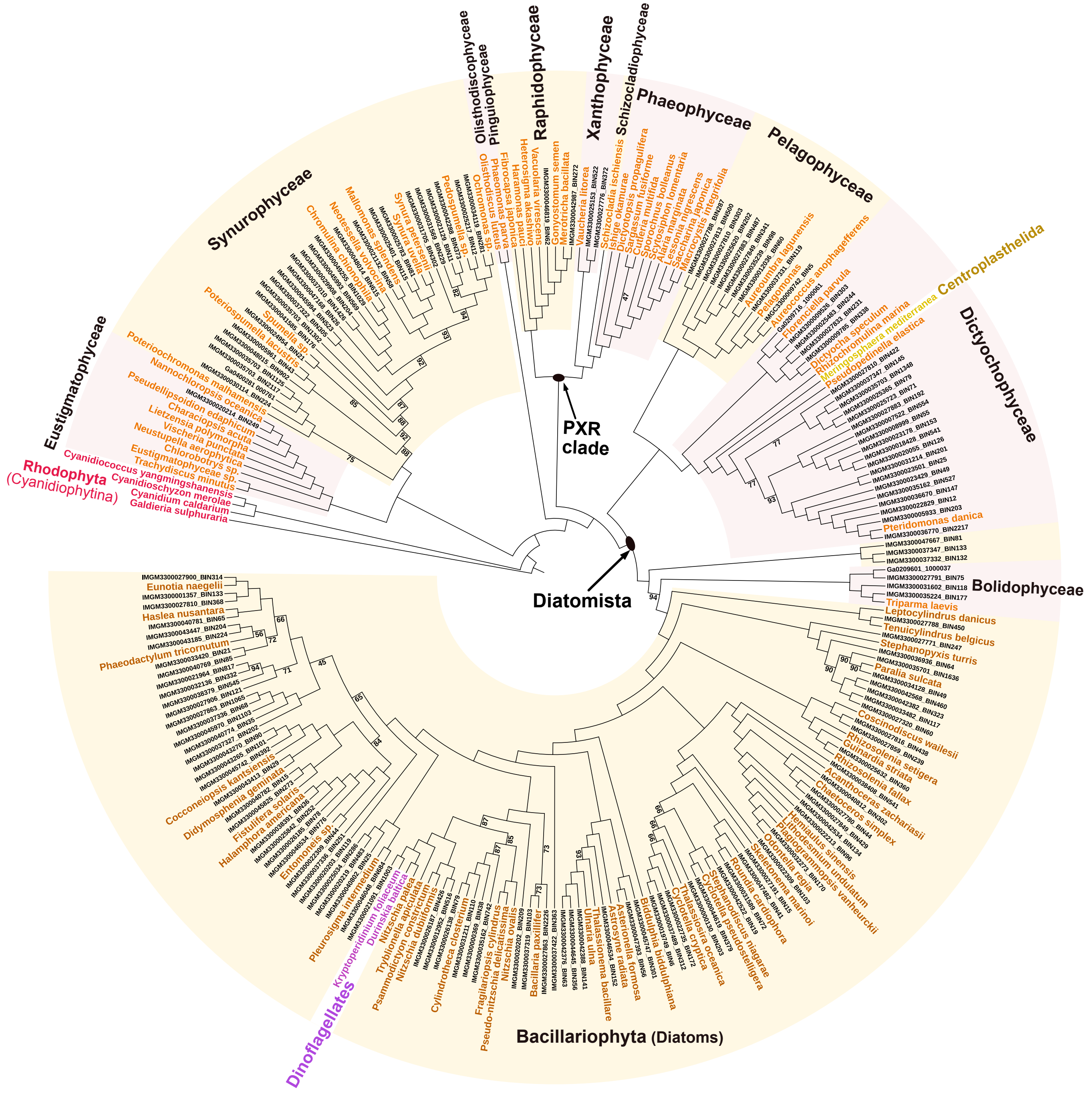


**Figure S7**. A cladogram depicting the phylogenetic relationships of secondary red algal plastids in Ochrophyta and other photosynthetic eukaryotes. The relationships were estimated using a minimum of 20% PLASTID54 markers using IQ-TREE and the LG+F+I+G4 substitution model. Red algal species (subphylum Cyanidiophytina) was used to root the tree. Only the branches with ultrafast bootstrap support values less than 95% are shown. Complete plastomes available in the NCBI RefSeq/GenBank are labeled with species name, while the ptMAGs are shown in black font. Major classes within Ochrophyta are distinguished with alternating pink or yellow shading and labeled. Plastids from Centroplasthelida and dinoflagellates are excluded from the shaded area. Reference ochrophyte species from the phylum Bacillariophyta are highlighted in darker orange compared to species from other ochrophytes.
