## Supplementary Text for "Global metagenomics reveals plastid diversity and unexplored algal lineages"

### RESULTS AND DISCUSSION

#### *Expanding the cyanobacterial taxonomy to explore plastid evolution*

Our initial phylogenetic reconstruction included 1,973 genomes/MAGs associated with Cyanobacteriota, using four Margulisbacteria species as an outgroup (Supplementary Fig. S1). This dataset was dereplicated to reduce representative cyanobacterial dataset without collapsing and mixing taxonomic orders. Through dereplication, the original dataset was reduced from 1,973 to 259 genomes/MAGs, comprising 16, 96, and 147 from the three clades Sericytochromatia, Vampirovibrionia, and Cyanobacteria, respectively. The phylogenetic relationships between the original and the dereplicated dataset were nearly identical, except the order FACHB-261 branched off prior to Gloeomargaritales in the dereplicated tree (Fig. 1A, Supplementary Fig. S1). Our phylogenetic analysis strongly supports monophyly of three major clades within Cyanobacteriota, placing Sericytochromatia at the base and the non-photosynthetic clade Vampirovibrionia as a sister to Cyanobacteria, consistent with published studies[1,2].

In terms of Cyanobacteriota taxon sampling, our study lacks species from the newly described GTDB order JACMPNO1 within Sericytochromatia but includes all other known orders. Within Vampirovibrionia, members formed two strongly supported monophyletic clades (UFBoot 100%) , includingVampirovibrionales and Gastranaerophilales as a sister to clade containing orders Obscuribacteriales, LMEP-6097, Caenarcaniphilales, and 2-02-FULL-35-15, consistent with other studies[1]. Gastranaerophilales represented the largest order within Vampirovibrionia with a total of 423 MAGs. Cyanobacteria was strongly supported as a monophyletic clade (UFBoot 100%). Gloeobacterales represented the most basal and early diverging order within Cyanobacteria, followed by the diversification of orders Thermostichales, Pseudanabaenales, and FACHB-261 with strong support (UFBoot >95%), consistent with previous studies[1,3]. Subsequently, the order Gloeomargaritales diversified from remaining cyanobacterial orders (moderate support, UFBoot 75%), followed by the order Thermosynechococcales (full support, UFBoot 100%). Our topology is consistent with a recent cyanobacterial phylogeny incorporating deep-branching lineages[3], although alternative relationships of Gloeomargaritales and Thermosynechococcales with other cyanobacterial orders have also been documented[1]. The remaining 14 photosynthetic cyanobacterial orders were grouped into two well supported clades: one including CAIUCS01, PCC7407, Elainellales, Phormidesmiales, CACIAM-69d, PCC-9006, Synechococcales, and PCC-9307, and the other containing orders SKYG9, GCF-001939115, FACHB-46, Neosynechococcales, Leptolyngbyales, and Cyanobacteriales (Fig. 1A). Cyanobacteriales represented the largest order, containing 558 genomes/MAGs in our dataset.

Our Cyanobacteria phylogeny included comprehensive taxon sampling from 21 orders and their relationships resembled the topology presented by Tan et al. (2024), except for the placement of the orders Elainellales and Phormidesmiales. Our topology strongly supports Elainellales as a sister to Phormidesmiales and together placed as a sister to PCC-6307 and Synechococcales, which is incongruent with the topology presented by Tan et al. (2024), where Elainellales was sister to Phormidesmiales and Cyanobacteriales.

We then utilized the dereplicated Cyanobacteriota representatives along with RefSeq NCBI plastomes to trace the origin of plastids, conducting several phylogenetic analyses while adjusting the phylogenetic markers (UNI56 and PLASTID54) and the ptMAGs dataset generated in our study. We noted that with an increasing presence of UNI56 markers (above 20%), more plastomes and ptMAGs representing Streptophyta and Chlorophyta were filtered out from the final alignment. This is mainly due to the large number of HMM profiles for ribosomal proteins, which constitute the majority of UNI56 markers. It is known that plastids in the red lineage have retained more plastid-encoded ribosomal proteins compared to those in the green lineage[4,5]. To mitigate this, we used PLASTID54 markers that included additional plastid-encoded photosynthesis-related proteins to retain plastomes /MAGs associated with the green lineage. Similarly, we observed that some ptMAGs, particularly those belonging to Streptophyta, shared high (>90%) average nucleotide identity (ANI) with RefSeq plastome as well mitochondrial genomes. For example, a ptMAG IMGM3300027552_BIN20 of a single contig of ~43 Kb, had 100% ANI and alignment fraction against *Arabidopsis thaliana* plastome (NC_000932), but also shared 92% ANI with a 91% alignment fraction against *Pulsatilla cernua* (from the family Ranunculaceae) mitochondrial genome (NC_068018). Plastid MAGs with higher ANI and alignment fraction against the plastome compared to the mitochondrial genome were included in our analysis. The presence of plastid sequences in mitochondrial genome due to intracellular and horizontal gene transfer are well documented in land plants[6–8], and likely explains the higher ANI shared with mitochondrial genomes. Furthermore, our initial phylogenetic analysis included Alveolata RefSeq plastomes, but due to their extremely long branch lengths, we excluded them and the associated ptMAGs from subsequently analyses (Supplementary Fig. S2).

In our phylogenetic analyses, ptMAGs and RefSeq plastomes together formed a strongly supported monophyletic group, positioned as a sister to cyanobacterial order Gloeomargaritales. This finding aligns with the observation that Archaeplastida plastids originated from deeply branching cyanobacterial lineage[9–11]. Whereas *Paulinella* plastids were placed within the recently diverged cyanobacterial order PCC-6307, following the diversification of *Synechococcus* sp. RSCF101. Our dereplicated dataset was much smaller, substantially reducing the taxa representing the order PCC-6307 from 498 to 28 taxa. Therefore, we inferred *Paulinella* plastid relationship by retaining all the original PCC-6307 taxa. Despite this, the phylogenetic position of *Paulinella* plastid within PCC-6307 remained unchanged (Supplementary Fig. S3), even with the expanded dataset. The *Synechococcus* sp. RSCF101 is distinct from *Synechococcus* sp. WH5701, which is considered closest extant relative of *Paulinella* plastid[12].

These two *Synechococcus* species belonged to separate phylogenetic clades within the PCC-6307 (Fig. 1D, Supplemental Fig. S3). Moreover, *Synechococcus* species WH5701 and RSCF101 exhibit noticeable differences in genomic features, including variation in genome sizes (3.04 Mb vs. 2.98 Mb, respectively) and gene counts (3403 vs. 3021 genes, respectively).

***Plastid MAGs and their relationships within Ochrophyta***

In their extensive phylogenomic analyses of stramenopiles, Derelle et al. (2016)[13] initially suggested two major divisions within Ochrophyta: Chrysista, which includes classes Xanthophyceae, Phaeophyceae, Rahidophyceae, Synchromophyceae, Synurophyceae, and Chrysophyceae, and Diatomista, which comprises the classes Pelagophyceae, Dictyochophyceae, Bolidophyceae, and the phylum Bacillariophyta (diatoms). Since then, several phylogenomic studies have aimed to delineate major ochrophyte lineages by sampling unrepresented taxa and improving phylogenetic resolution. However, the current understanding of Ochrophta relationships remains inconsistent due to incomparable taxon sampling, different markers (nuclear and organellar), and phylogenetic methods used[14–18]. While the relationships among the major classes within Diatomista are well resolved, incongruent relationships among the classes within Chrysista are prevalent.

In our study, most novel ptMAGs recovered belonged to Ochrophyta (n=155, Fig. 2). We reconstructed phylogenies by combining the novel ptMAGs and reference ochrophyte plastomes to assess the placement of novel ptMAGs within ochrophytes. Our phylogenetic analyses strongly support the monophyly of Ochrophyta but with inconsistent placement of some classes belonging to Chrysista (Fig. 2, Supplementary Figs. S5-S7). Our phylogenetic reconstruction, which included only Ochrophyta plastomes and ptMAGs consistently recovered Chrysista as a paraphyletic group (Supplementary Figs. S5-S7). To avoid the influence of novel ptMAGs, we first reconstructed the phylogeny using only 213 reference ochrophyte plastomes (Supplementary Fig. S6). The reference ochrophytes phylogeny recovered two strongly supported clades: one with Synurophyceae and Eustigmatophyceae and another clade with remaining all other ochrophytes. Olisthodiscophyceae and Pinguiophyceae, each represented by a single species, formed a moderated supported clade (UFBoot 90%), and diverged early as sister to PXR and Diatomista. The PXR clade was strongly supported as a monophyletic group (UFBoot 100%) and was recovered as a sister to Diatomista with relatively low support (UFBoot 78%). Within Diatomista, Pelagophyceae and Dictyochophyceae formed a moderately supported clade as a sister to Bolidophyceae and Bacillariophyta (UFBoot 93%). When we added novel ptMAGs associated with Ochrophyta on the reference dataset, an identical topology was recovered with much higher bootstrap support (Supplementary Fig. S7). This topology lacks the dichotomy between Chrysista and Diatomista and is incongruent with the current phylogenomic studies[14,17,18], likely due to the phylogenetic makers. We selected plastid phylogenetic markers commonly present in red and green lineages with the broad aim to understand plastid evolution across diverse photosynthetic eukaryotes. To resolve evolutionary relationships with

Ochrophyta, lineage-specific phylogenetically informative markers, robust assessment of phylogenetic methods, and comprehensive taxon sampling are required[14,18]. Despite the incongruency in relationship between Ochrophyta classes, the placement of novel ptMAGs within the classes was always consistent. Therefore, we used the inferred topology to describe the placement of Ochrophyta-related novel ptMAGs (Supplementary Fig. S7).

Among 155 Ochrophyta-related novel ptMAGs, most (n=122) belonged to Diatomista, with the remaining 33 placed within Chrysista (Supplementary Fig. S7). Within Chrysista, the largest number of ptMAGs (n=27) were within Synurophyceae, with size ranging from ~19 Kb to ~132 Kb. The remaining ptMAGs within Chrysista were confined to Raphidophyceae (n=3), Xanthophyceae (n=2), and a single novel ptMAG of ~110 Kb within Eustigmatophyceae, placed as a sister to *Nannochloropsis* sp., and shared near identical synteny with *Nannochloropsis* plastome (result not shown). Within Diatomista, novel ptMAGs were distributed across all major clades. Pelagophyceae contained ten novel ptMAGs, size ranging from ~47 Kb to ~138 Kb. Similarly, 24 novel ptMAGs, ranging in size from ~21 Kb to ~110 kb were recovered within Dictyochrophyceae. We also recovered the plastid of the centroplasthelida species *Meringosphaera mediterranea* (a Haptista) between *Rhizochromulina marina* and *Pseudopedinella elastica* plastids. Plastids in *Meringosphaera* are known to be derived through kleptoplasty, a process where plastids of engulfed prey are temporarily retained. Phylogenetic analyses using 16S rDNA and plastid markers have placed them within Dictyochrophyceae[19], consistent with our phylogenetic placement.

We consistently recovered a clade containing only ptMAGs prior to the diversification of Bolidophyceae and Bacillariophyta (Supplementary Fig. S7). The clade included three novel ptMAGs, ranging in size from ~46 Kb to ~49 Kb, which lacked reference plastomes, and were strongly supported (UFBoot 100%) as sister to Bolidophyceae and Bacillariophyta. The class Bolidophyceae is represented by a single complete plastome of *Triparma laevis*[20]. We identified a clade that included four additional ptMAGs, ranging from ~104 Kb to ~106 Kb, along with *T. laevis*. These metagenomes are likely to be near complete, as their sizes are comparable to the ~112 Kb *T. laevis* plastome (excluding a copy of inverted repeats). Within Bolidophyceae, *T. laevis* plastome, together with four novel ptMAGs formed two distinct clades within Bolidophyceae. One contained the *T. laevis* plastome along with two ptMAGs, IMGM3300031602_BIN118, and IMGM3300035224_BIN177, and the other clade with two metagenomes IMGM3300025666_Ga0209601 and IMGM3300027791_BIN75. Interestingly, two copies of *psbA* gene, which is suggested to be a unique feature of the *T. laevis* plastome, were also found in two ptMAGs IMGM3300031602_BIN118, and IMGM3300035224_BIN177. In contrast, only a single copy of *psbA* gene was present in the sister clade containing only ptMAGs. In IMGM3300035224_BIN177, a copy of *psbA* was much smaller, about 265 bp compared to the full-length gene of 1083 bp and was located at the end of a contig, likely representing a partial *psbA* copy due to incomplete assembly. The two copies of *psbA* in *T. laevis* were attributed to gene duplication after the diversification of Bolidophyceae[20]. However, the absence of the duplicated copy in the sister clade within Bolidophyceae suggests the duplication event was confined to a specific clade within Bolidophyceae. Alternatively, the gene duplication may have occurred early during the speciation of Bolidophyceae, with the sister clade lacking the duplicate *psbA* due to incomplete assembly.

Bacillariophyta was strongly supported as a monophyletic clade and included the largest number of novel ptMAGs (n=81), with assembly sizes ranging from ~20 Kb to ~116 Kb. The relationship among Bacillariophyta families recovered in our phylogenetic reconstructions were consistent with the current comprehensive phylogenomic study using concatenated plastid genes of reference diatom species[21]. The extensive collection of novel ptMAGs associated with Bacillariophyta generated in this study will serve as a valuable resource for future research, significantly enhancing phylogenetic resolution and facilitating comparative analyses within this group.

Lastly, we identified a single novel ptMAG (IMGM3300021091_BIN1003) that is strongly supported (UFBoot 100%) as a sister to plastome of dinoflagellate species *Kryptoperidinium foliaceum* (Supplementary Fig. S7). The ptMAG is represented by two contigs with a total length of ~108 Kb and retains nearly all protein coding genes present in *K. foliaceum* plastome except for *ccsA* (Cytochrome C biogenesis), *tyrC* (tyrosine recombinase), and *serC1/C2* (serine recombinases), likely due to incomplete assembly. Plastids of *K. foliaceum* and the closely related dinoflagellate species *Durinskia baltica* are thought to be derived from a diatom through tertiary endosymbiosis and are collectively referred to as dinotoms[22]. Early phylogenetic studies using single gene markers and rDNA sequences suggested dinotom plastids were closely related to plastids of *Nitzschia* species[23,24]. Due to the lack of complete plastomes from *Nitzschia* species at the time, dinotom plastomes were compared with those of the free-living diatom species *Phaeodactylum tricornutum*[22]. With the access to multiple *Nitzschia* plastomes, *D. baltica* plastome was found to be highly similar (92% pairwise identity) to the *Nitzschia palea* plastome, sharing identical gene content and synteny[25]. However, monophyletic dinotoms were found to retain plastids of multiple *Nitszchia* or other diatom species through independent acquisition and replacement of the endosymbiont[26]. This complicates the understanding of tertiary endosymbiosis in dinotoms, making it challenging to pinpoint the source of endosymbiont plastids.

In our phylogenetic reconstruction, plastids from two dinotoms (*K. foliaceum* and *D. baltica*) formed a strongly supported (UFBoot 100%) monophyletic group with *Nitzschia palea, Tryblionella apiculata,* and the ptMAG IMGM3300021091_BIN1003 (Supplementary Fig. S7). Within the group, *T. apiculata* branched off early at the base, with *K. foliaceum* and IMGM3300021091_BIN1003 grouping as a sister to *D. baltica* and *N. palea*. It is likely that *K. foliaceum* may have retained the plastid of the diatom *T. apiculata,* whereas in *D. baltica,* their plastids were replaced with the endosymbiont *N. palea.* The plastome size (excluding a copy of inverted repeats) of *K. foliaceum* (134,409 bp) was comparable to that of diatom *T. apiculata* (129,773 bp) and shares all protein coding genes except a copy of putative serine recombinase (*serC1*). Although *T. apiculata* plastome lacks annotated tyrosine recombinase (*tyrC)* found in *K. foliaceum,* it contains a putative integrase/recombinase gene annotated as *xerC*[27]. The XerC is 101 amino acids long, substantially shorter (due to premature stop codon) compared to TyrC of 312 aa, and shares 52% pairwise aa identity, with both containing an integrase catalytic domain (IPR002104). The pairwise nucleotide identity between these two genes are much higher (63%) with the upstream region of *xerC* gene sharing 59% nucleotide identity with the 5' region of the *tyrC* gene. The nearly identical gene content between the dinotom *K. foliaceum* and the diatom *T. apiculata* suggests that *K. foliaceum* may have retained the plastid of the diatom *Tryblionella apiculata,* whereas in *D. baltica,* their plastids were replaced with the endosymbiont *N. palea.*

Our phylogenetic reconstruction Ochrophyta, incorporating novel ptMAGs and RefSeq plastomes, supports a monophyletic origin of Ochrophyta plastids but reveals inconsistencies in placements for some Chrysista classes. Subsequent studies with lineage-specific markers and robust phylogenetic methods will likely solve these relationships, and the novel ptMAGs will be resourceful. Despite these inconsistencies, the novel ptMAGs consistently fit within their respective classes, with most belonging to Diatomista. We identified many novel clades represented solely by ptMAGs but lacking reference plastomes in including Synuophyceae, Pelagophyceae, Dictyopchophyceae, Bolidophyceae and the phylum Bacillariophyta. Among them, an early branching clade sister to Bolidophyceae and Bacillariophyta, likely represents a novel lineage with essential implication on our understanding of plastid evolution in diatoms. Furthermore, novel ptMAGs closely related to the dinotoms like *K. foliaceum* and *D. baltica*, show close relationships with specific diatom species and provide resources to investigate evolution of complex plastids in dinoflagellates.

### PLASTID MAG CONSORTIUM

Katherine McMahon, University of Wisconsin-Madison; Ramunas Stepanauskas, Bigelow Laboratory for Ocean Sciences; Alison Buchan, University of Tennessee-Knoxville; Thomas Mock, University of East Anglia; Kirsten Fisher, California State University; Joan Slonczewski, Kenyon College; and Luce Ward, Smith College.
